# Dbp1 is a low performance paralog of RNA helicase Ded1 that drives impaired translation and heat stress response

**DOI:** 10.1101/2024.01.12.575095

**Authors:** Emily N Powers, Naohiro Kuwayama, Camila Sousa, Kendra Reynaud, Marko Jovanovic, Nicholas T Ingolia, Gloria A Brar

## Abstract

Ded1 and Dbp1 are paralogous conserved RNA helicases that enable translation initiation in yeast. Ded1 has been heavily studied but the role of Dbp1 is poorly understood. We find that the expression of these two helicases is controlled in an inverse and condition-specific manner. In meiosis and other long-term starvation states, Dbp1 expression is upregulated and Ded1 is downregulated, whereas in mitotic cells, Dbp1 expression is extremely low. Inserting the *DBP1* ORF in place of the *DED1* ORF cannot replace the function of Ded1 in supporting translation, partly due to inefficient mitotic translation of the *DBP1* mRNA, dependent on features of its ORF sequence but independent of codon optimality. Global measurements of translation rates and 5’ leader translation, activity of mRNA-tethered helicases, ribosome association, and low temperature growth assays show that—even at matched protein levels—Ded1 is more effective than Dbp1 at activating translation, especially for mRNAs with structured 5’ leaders. Ded1 supports halting of translation and cell growth in response to heat stress, but Dbp1 lacks this function, as well. These functional differences in the ability to efficiently mediate translation activation and braking can be ascribed to the divergent, disordered N- and C-terminal regions of these two helicases. Altogether, our data show that Dbp1 is a “low performance” version of Ded1 that cells employ in place of Ded1 under long-term conditions of nutrient deficiency.

## Introduction

DEAD-box ATPases, named for an amino acid motif involved in their catalytic activity, are a highly conserved and abundant class of proteins with critical roles in the regulation of RNA (Bohnsack et al., 2023; Putnam and Jankowsky, 2013; Sharma and Jankowsky, 2014; Weis and Hondele, 2022). Their best characterized role is as helicases, functioning as non-processive remodelers of RNA secondary structure, which aids them in functions including mRNA decay, ribosome biogenesis, mRNA export, and translation initiation. They are superfamily 2 nucleic acid helicases, sharing a conserved helicase core containing two RecA domains, flanked in most cases by diverse, protein-specific disordered extensions on the N- and C-termini (Weis and Hondele, 2022).

Translation initiation—the process by which start codons within mRNAs are chosen and decoded by the ribosome—requires two DEAD box proteins in yeast, Ded1 (DDX3 in mammals) and eIF4A (Tif1/2 in yeast; DDX2 in mammals; (Andreou and Klostermeier, 2013; Sharma and Jankowsky, 2014; Weis and Hondele, 2022)). This process occurs predominantly by the scanning mechanism, beginning with binding at an mRNA 5’ end by the 43S preinitiation complex (PIC)—including the 40S ribosomal subunit, initiator methionine tRNA, GTP, and several initiation factors—directed by 5’ cap-bound eIF4G, eIF4E, and eIF4A (referred to together as eIF4F) to form the 48S PIC. 48S complexes scan the mRNA in a 5’ to 3’ direction, traversing the 5’ leader region until a suitable start codon is found (Hinnebusch, 2014). Ideal start codons are AUGs with optimal (or “Kozak”) sequence context, but AUGs in poor sequence context or non-AUGs are chosen in some instances in mutant backgrounds, and more rarely within specific transcripts in wild-type cells (Brar, 2016; Hinnebusch, 2014, 2011; Kozak, 1986). Scanning is enabled by the RNA unwinding activity of eIF4A, as well as Ded1, which help to resolve secondary structures that impede progress of the PIC (Gao et al., 2016; Guenther et al., 2018a; Gulay et al., 2020; Gupta et al., 2018; Sen et al., 2015; Zhou et al., 2023). Based on this important role, both eIF4A and Ded1 are essential to bulk translation and cell viability.

eIF4A appears to be important for translation initiation on all mRNAs with little specificity, whereas Ded1 has been shown to have particular importance for translation initiation at start codons downstream of highly structured 5’ leaders (Firczuk et al., 2013; Hilliker et al., 2011; Sen et al., 2015; Zhou et al., 2023), a role shared by its human homolog DDX3 (Calviello et al., 2021). Diminished function of Ded1 leads to lower translation levels and reduced growth rates (Chuang et al., 1997; Sen et al., 2015). Ded1 associates with PICs through its interactions with each of the components of eIF4F, and is important for efficient 48S PIC formation (Aryanpur et al., 2019; Gao et al., 2016; Gulay et al., 2020; Hilliker et al., 2011; Senissar et al., 2014; Zhou et al., 2023). Examination of translation initiation in cells deficient for Ded1 revealed spurious translation initiation at a subset of near-cognate codons within 5’ leaders, and particularly at sites upstream of structured regions of mRNA (Aryanpur et al., 2017; Guenther et al., 2018a). Paradoxically, Ded1 overexpression is also associated with repression of translation through stress granule formation, and this function is also seen under specific stress conditions, including heat stress, in the absence of its overexpression (Aryanpur et al., 2017; Beckham et al., 2008; Hilliker, 2012; Hilliker et al., 2011; Iserman et al., 2020). The ability to phase separate is inherent to Ded1 itself, as purified protein forms condensates in vitro (Weis and Hondele, 2022).

Dbp1 is a paralog of Ded1 (also homologous to DDX3 in mammals) with a core amino acid sequence that is 82.6% identical to Ded1 and more divergent N- and C- termini sequences (Figure 1A; (Banroques et al., 2011; Jamieson and Beggs, 1991; Sen et al., 2019)). Dbp1 is nonessential, very lowly expressed under standard laboratory growth conditions, and it can support growth in cells lacking Ded1 but only when expressed at very high levels (Berthelot et al., 2004; Jamieson and Beggs, 1991; Sen et al., 2019). Ded1 has intrinsically disordered N- and C-termini that contain its eIF4F binding sites and mutations of these regions convey loss of function phenotypes *in vivo* (Banroques et al., 2011; Floor et al., 2016; Gao et al., 2016; Gulay et al., 2020; Hilliker et al., 2011; Senissar et al., 2014). Whether equivalent sites exist within Dbp1 termini has not been determined. In vitro comparison of Ded1 and Dbp1 activity has shown that Dbp1 has lower RNA duplex unwinding activity and lower RNA-stimulated ATPase activity than Ded1 (Banroques et al., 2011), but greater ability to accelerate 48S PIC formation on structured 5’ leaders (Gupta et al., 2018; Sen et al., 2019).

**Figure 1:**
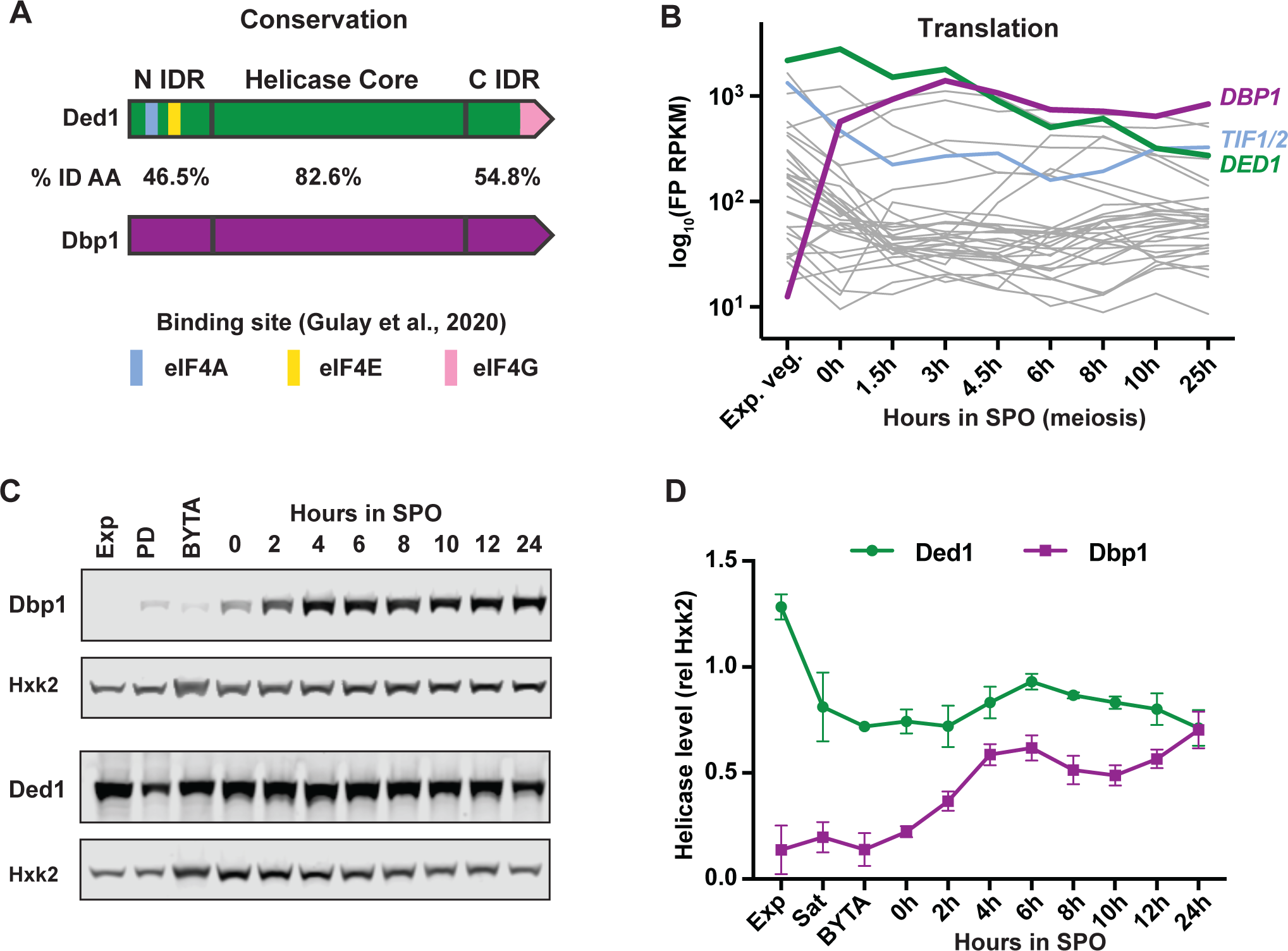
Dbp1 is upregulated and Ded1 is downregulated during meiosis, relative to mitotic growth. (A) Conservation of Dbp1 and Ded1 as shown by amino acid percent identity (AA % ID) of the N-terminal, core, and C-terminal regions, as defined by Banroques et al., 2011. Location of initiation factor binding regions of Ded1 from (Aryanpur et al., 2019; Gulay et al., 2020) are indicated. (B) Translation levels, as assessed by ribosome footprint (FP) quantification by RPKM of all yeast RNA helicases. Data are from (Cheng et al., 2018b) and represent the average of two biological replicates. Translation of known and putative RNA helicases shown in grey, translation initiation related helicases *DBP1*, *DED1*, and *TIF1/2* are labeled. Note that “sat” refers to saturated cells, which are in the post-diauxic growth phase (C) Western blots and quantification (D) demonstrate an accumulation of Dbp1 and a reduction in Ded1 levels over the course of meiosis. Data represent the average of three biological replicates with error bars reflecting standard deviation.

Dbp1 helicase can perform similar functions to Ded1 but it is unclear whether Dbp1 has any unique functional roles or characteristics (Berthelot et al., 2004; Jamieson and Beggs, 1991; Powers et al., 2022; Sen et al., 2019; Weis and Hondele, 2022). In vivo analysis of Dbp1 function has been challenging due to a dearth of conditions in which it is known to be expressed and an unexpected side-effect associated with its deletion via cassette-based genomic replacement. We recently showed that this genome editing strategy, which was used to make all published yeast strains deleted for the *DBP1* locus (Banroques et al., 2011, 2010, 2008; Berthelot et al., 2004; Sen et al., 2019), causes strong translational down-regulation of *DBP1*’s neighboring ORF, encoding the mitochondrial ribosomal protein Mrp51. This is a result of the inherent bidirectionality of the minimal promoters used in expression cassettes (Powers et al., 2022). This off-target effect results in downregulation of cytosolic translation and was responsible for all mutant phenotypes observed in cells deleted for *DBP1* via cassette replacement (Powers et al., 2022).

One condition in which Dbp1 is highly expressed is meiosis, a highly specialized cell differentiation program that produces haploid gametes from diploid cells. Induction into meiosis is driven by a change in nutrient state in budding yeast, as cells are shifted from a condition in which there are abundant nitrogen and fermentable carbon sources, to one in which only non-fermentable carbon sources are available and nitrogen is low (reviewed in (Broach, 2012; Neiman, 2011; Plank, 2022)). Timely induction of this differentiation program is crucial for population survival when cells experience nutrient deprivation conditions, as the resultant spores are highly fortified and resistant to many stressors (reviewed in (Broach, 2012; Neiman, 2011; Plank, 2022). Furthermore, if a growing yeast population depletes its available nutrients and continues to divide mitotically, cells will eventually starve and perish unless growth is quelled or differentiation is stimulated.

The discovery that any direct Dbp1-dependent mutant phenotypes found in previous studies of *dbp1Δ* cells were likely to be masked by off-target effects, together with strong upregulation of Dbp1 expression in specific conditions including meiosis, motivated us to investigate its function. Our studies were designed to investigate Dbp1 and Ded1 roles in vivo, at endogenous and equivalent levels, given the known concentration-dependence of Ded1’s translation activating and repressing roles. We also avoided use of temperature-sensitive alleles to prevent confounding effects of temperature on translation. Our work uncovered extremely tight regulation of expression for both helicases, in part through translational regulation of the transcripts that encode them. Dbp1 and Ded1 protein synthesis is inversely correlated, and unexpectedly, the translation level of *DBP1* depends on its ORF sequence but not its codon optimality.

We show that replacement of Ded1 with equivalent levels of Dbp1 in exponentially growing mitotic cells leads to enhanced translation at near-cognate codons and poorer translation of a subset of mRNAs with highly structured 5’ UTRs, arguing that Dbp1 is less effective at stimulating translation than Ded1. Supporting this model, compared to Ded1, Dbp1 associates less efficiently with translating ribosomes, is less effective at activating translation when tethered to a model transcript, and fails to support cell growth at low temperatures when mRNA structure would be expected to be more stable. Cells expressing Dbp1 in place of Ded1 are also unable to halt growth at high temperatures, a role attributed to translation inhibition through stress granule formation. Our data show that its divergent and disordered termini are responsible for both Dbp1’s deficiencies in translation activation and inhibition, relative to Ded1. In conditions of rapid growth, Ded1 is the sole helicase expressed, allowing cells to maximally activate translation and rapidly halt translation in response to changes in external conditions. Dbp1 expression is upregulated and Ded1 downregulated specifically in several conditions of long-term nutritional stress, suggesting the that a “low performance” DEAD-box helicase like Dbp1, with less ability to stimulate *and* rapidly halt translation, is preferred under conditions in which rapid responses to changes in environment are not advantageous.

## Results

### Dbp1 and Ded1 expression are inversely correlated

Ded1 is highly expressed during mitotic exponential growth but it was reported that its translation decreases during meiosis. In contrast, its paralog, Dbp1, is translated at a level that is 69-fold higher in meiotic cells relative to mitotic cells (Figure 1B; (Brar et al., 2012; Guenther et al., 2018)). Dbp1 is unusual in this respect, as one of only four translation-associated RNA helicases that increase in translation during meiosis, and the one that increases with the highest magnitude of this set. Most behave similarly to eIF4A (encoded by *TIF1* and *TIF2*), decreasing initially and remaining low, consistent with the role of many RNA helicases in facilitating ribosome biogenesis and a decrease in ribosome biogenesis and bulk translation early in meiosis (Brar et al., 2012; Martin et al., 2013; Rodríguez-Galán et al., 2013). Dbp1 and Ded1 protein abundances follows a similar trend as their translation, with Ded1 decreasing to roughly half its mitotic levels in mid-meiosis (Figures 1C, 1D). This decrease was previously observed—but was even more pronounced—in a study that used meiotic cells that were hyper-synchronized (Carlile and Amon, 2008; Guenther et al., 2018a). Dbp1, by comparison, is typically undetectable by western blotting in mitotic cells (Figure 1C). Mitotic *DBP1* mRNA abundance (as assessed by mRNA-seq) was also very low, at ∼2 reads per kilobase million (RPKM; (Brar et al., 2012; Cheng et al., 2018b)), which is equivalent to *SPO11* and *DMC1*, two well-studied meiotic transcripts known to be restricted from mitotic expression. *DBP1* mRNA and translation are induced in meiosis, with its translation eventually increasing to levels that are comparable to those of Ded1 in late meiosis ((Brar et al., 2012; Cheng et al., 2018b); Figures 1C, 1D). This regulation occurs through increased mRNA abundance via transcriptional induction, rather than reduced mRNA degradation, as a mitotic nascent transcript sequencing study did not observe *DBP1* mRNA to be transcribed in mitotic cells (Churchman and Weissman, 2011).

A microarray study that examined many growth and stress conditions in budding yeast identified several non-meiotic conditions in which *DBP1* mRNA was expressed, including “post-diauxic” growth following exhaustion of a fermentable carbon source, stationary phase (an extended cell cycle arrest driven by nutrient exhaustion), and long-term nitrogen deprivation (Figure 1SA;(Gasch et al., 2000)). Notably, in all cases in which Dbp1 expression is induced, Ded1 expression decreases. Dbp1 expression is not induced in response to the majority of characterized cellular stresses, including amino acid starvation, long-term temperature shifts or temperature shock, osmotic, oxidative, or unfolded protein stresses (Gasch et al., 2000). Increased Dbp1 expression and reduced Ded1 expression thus appear to be associated with long-term states of cellular maintenance in specific limiting nutrient conditions.

We wondered if the inverse relationship between the expression of these two helicases led to a change in helicase activity. In particular, we noted that an increase in translation within 5’ UTRs, particularly at non-AUG codons, occurs when Ded1 function is compromised, attributed to inefficient disruption of 5’ structures during PIC scanning (Guenther et al., 2018b; Kearse and Wilusz, 2017; Wang et al., 2022). Meiotic cells also display increased 5’ leader translation and non-AUG initiation (Figure S1B;(Brar et al., 2012; Guenther et al., 2018a; Hollerer et al., 2021)). We reasoned that if Dbp1 could not fully substitute for Ded1 function in efficient 5’ leader scanning and subsequently stringent translation initiation, the increase in Dbp1 and decrease in Ded1 expression seen during meiosis could contribute to meiotic translation within 5’ leaders (Figure 1SB).

### Genomic replacement of DED1 with DBP1 reveals their ORF-dependent differential translation

Dbp1 can substitute for Ded1’s function in supporting cell growth when expressed from a high copy plasmid (Jamieson and Beggs, 1991; Sen et al., 2019), but these conditions would be expected to result in massive overexpression of Dbp1 relative to endogenous levels of either helicase. To test whether Dbp1 helicase had characteristics unique from Ded1, we asked whether equivalent levels of Dbp1 could rescue mitotic exponential growth in cells lacking Ded1 helicase. We chose these conditions because Ded1 is essential and has defined functions in supporting mitotic growth (Sen et al., 2015; Struhl, 1985). We used CRISPR/Cas9 to create an un-marked deletion of *DED1* (*ded1Δ).* In diploid cells lacking the *DED1* open reading frame (ORF) at its endogenous locus, we integrated a construct homozygously at the *LEU2* locus that contained either the *DED1* or *DBP1* ORF under control of the *DED1* promoter, 5’ leader, 3’ UTRs, and terminator (Figure 2A). Untagged Ded1 produced by this strategy supported normal mitotic exponential growth, resulting in the typical doubling time of ∼90 minutes. (Figures 2B, 2C). However, untagged Dbp1 did not, causing cell doubling time to be increased by roughly 20% (Figures 2B, 2C). This reduced growth rate was associated with decreased bulk translation in mitotic cells expressing Dbp1 in place of Ded1, as assessed by measurement of ^35^S amino acid incorporation and polysome gradient analysis (Figures 2D, 2E). This suggested that Dbp1 was less effective at supporting translation than Ded1 in mitotic cells.

**Figure 2:**
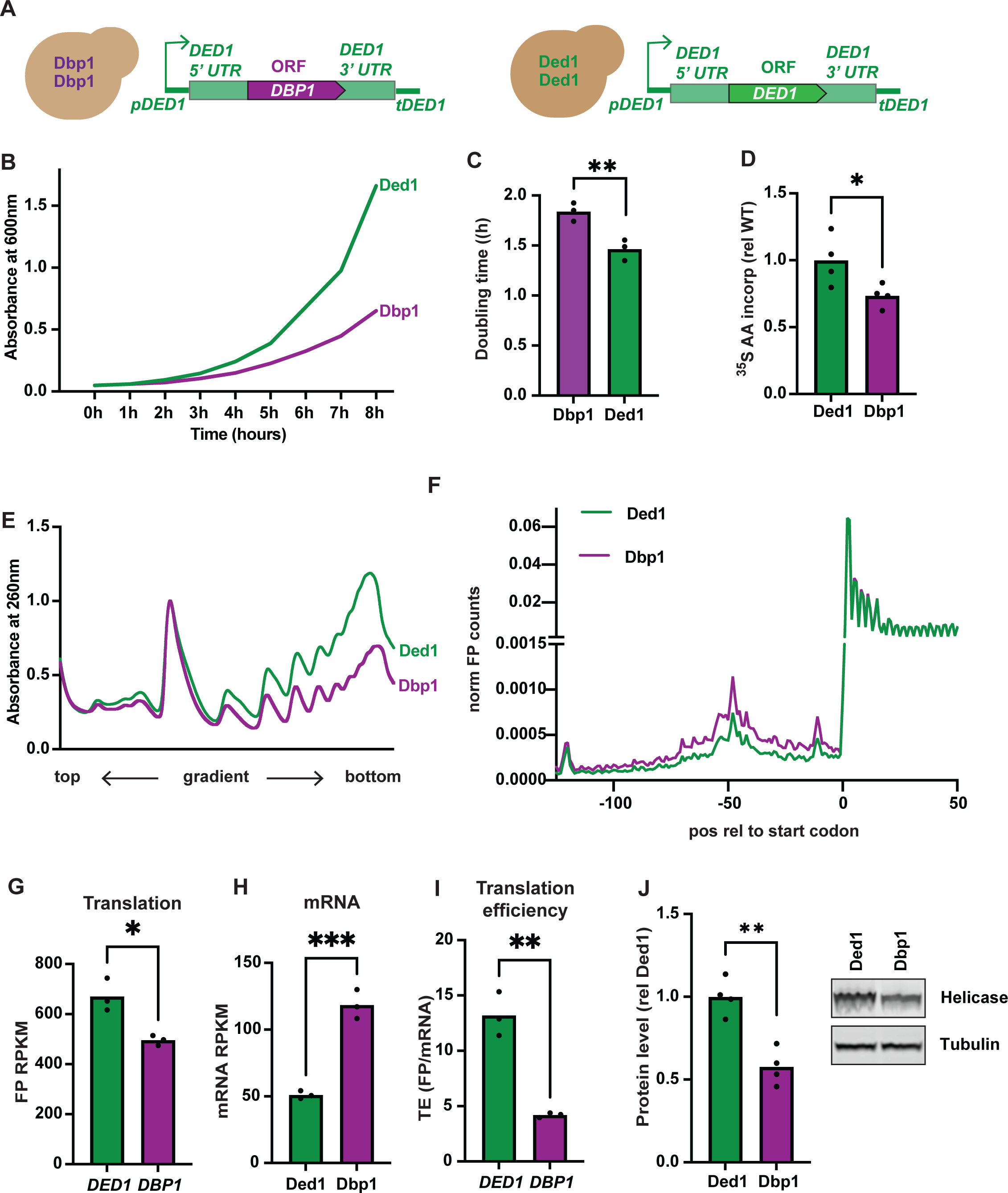
DBP1 ORF cannot substitute for DED1 ORF in supporting robust mitotic growth or translation. (A) Schematic of constructs integrated in single copy at the *LEU2* locus. Diploid cells were deleted for the *DED1* ORF at its endogenous locus and either the *DBP1* or *DED1* ORF is flanked by *DED1* regulatory regions and integrated at *LEU2*. (B) Growth curves of Ded1- or Dbp1- expressing cells under mitotic exponential conditions (rich media; YEPD). Three biological replicates were analyzed, one representative trace is shown. (C) Calculated doubling time for all replicates. (D) Rates of total translation of cells expressing Dbp1 or Ded1 under mitotic exponential growth conditions as determined by radioactive amino acid incorporation assays. Data represent measurements from four biological replicates, ^35^S amino acid incorporation rates for Ded1 and Dbp1 expressing strains are shown relative to wild-type rates of incorporation. Statistical significance was determined by a two-tailed P value < .05 resulting from unpaired t test. (E) Polysome profiles of Ded1 or Dbp1 expressing cells under mitotic exponential growth conditions matched to (B). Three biological replicates were analyzed, one representative trace is shown. (F). Metagene plots demonstrating ribosome FP occupancy across all genes relative to their canonical AUG start codon. The sum of FP counts at each position was normalized to the total mapped FP counts across the metagene plot for each sample. Three biological replicates were analyzed and the average normalized FP counts are shown. (G-I) translation (G), mRNA (H) and translation efficiency (TE; I) of *DBP1* relative to *DED1* in cells under mitotic exponential growth conditions. Data from three biological replicates are presented and statistical significance indicates a two-tailed P value < .05 = *, P < .01 = **, and P < .001 = *** as determined by unpaired t test. (J) Levels of Ded1 and Dbp1 helicase under mitotic exponential growth conditions as determined by western blotting. Data from four biological replicates are shown, significance indicates a two-tailed P value < .05 as determined by unpaired t test.

The bulk translation defect seen when Ded1 is replaced by Dbp1 could be transcript-specific, meaning that Ded1 is required to promote the efficient translation of specific transcripts that support rapid growth, and ultimately high translation. Reduced bulk translation in this case would be an indirect rather than direct result of the helicase type present. Alternatively, Dbp1 could have a lower capacity than Ded1 to stimulate translation initiation on all transcripts, thus supporting lower bulk translation and slower growth than mitotic cells expressing Ded1. To distinguish between these possibilities, we measured translation efficiencies (TEs) sgenome-wide in these strains by performing ribosome profiling and mRNA-seq in triplicate on Ded1- versus Dbp1*-*expressing cells (Figure 2A; S2). Data were reproducible, as assessed by replicate analysis (Figure S2A). Increased ribosome density in the region 100 nucleotides (nts) upstream of annotated ORF start codons was observed in cells expressing Dbp1 compared to Ded1 (Figure 2F), suggesting that Dbp1 could not stimulate 5’ leader scanning and robust initiation to the degree that Ded1 does. Additionally, many genes displayed differences in mRNA, translation rates, and translation efficiencies (TEs; ribosome footprint RPKM/mRNA RPKM) between Dbp1- and Ded1-expressing cells (Figure S2B-G).

Interestingly, despite matched regulatory regions driving Dbp1 and Ded1, translation of Dbp1 was significantly lower than translation of Ded1 (Figure 2G, S2H). *DBP1* mRNA levels in Dbp1-expressing cells were 2.4-fold higher than that of *DED1* in Ded1-expressing cells (Figure 2H), resulting in a TE for *DBP1* that was less than half of that observed for *DED1* (Figure 2I). As expected based on translation rates (Figure 2G), Dbp1 protein was present at a significantly lower abundance than Ded1 (Figure 2J), as assessed by western blotting of strains expressing C-terminally 3V5-tagged helicases and a second set of strains with internal 3V5 tags for both helicases (Figure 3A; epitope tag inserted between N-terminal 220 and 221 amino acids of Dbp1 and between N-terminal 213 and 214 amino acids of Ded1). We confirmed using RT-qPCR and western blotting (Figure 3SA; Figure 3A), that mRNA levels were increased and protein levels decreased, respectively, for Dbp1 compared to Ded1 when either was expressed from the *DED1* regulatory regions, resulting in a protein to mRNA ratio that was almost 3-fold higher for *DED1* than *DBP1* (Figure 3A).

**Figure 3:**
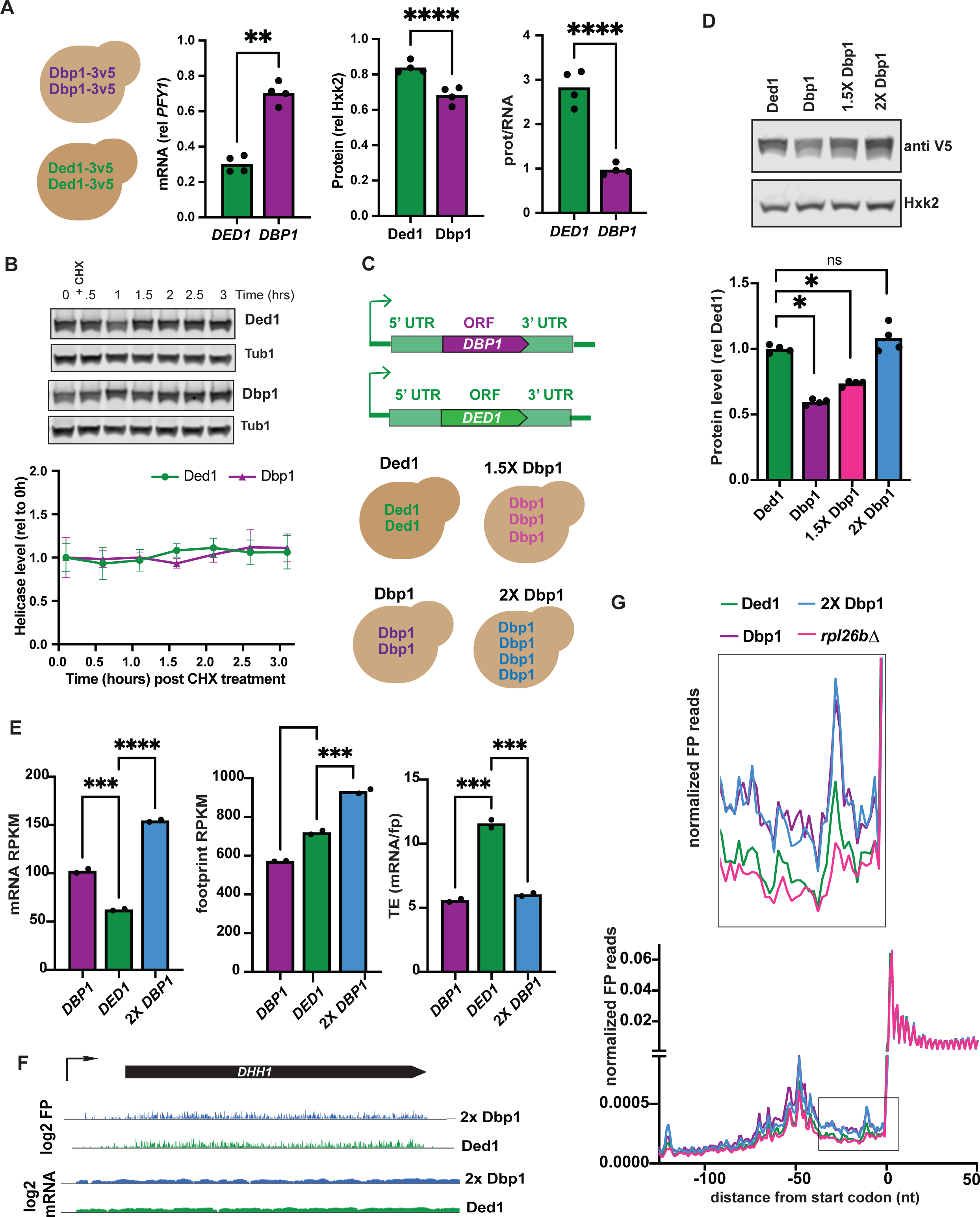
DBP1 ORF is less efficiently translated than DED1 ORF. (A) Top: Schematic demonstrating the genetic background of diploid strains expressing homozygous single copy integrations of tagged helicase alleles from the *DED1* regulatory sequences. Bottom: Comparisons of *DED1/DBP1* mRNA levels relative to *PFY1* as determined by RT-qPCR, protein levels relative to Tub1 as determined by western blotting , and the resulting protein/mRNA ratio for the *DBP1* and *DED1* encoding transcripts. Data represent four biological replicates and statistical significance was determined by a statistical significance indicates a two-tailed P value < .05 = *, P < .01 = **, P < .001 = ***, and P < .0001 = ****. (B) Western blots and quantifications demonstrating Dbp1 or Ded1 levels following translation shut-off by addition of cycloheximide. Data points represent the average of three biological replicates and error bars represent standard deviation. (C) Schematic of single copy genomic integrations as defined in Figure 3SC. (D) Quantification of Dbp1 protein levels as compared to Ded1 in single copy replacement strain (Dbp1), heterozygous extra copy strain (1.5X Dbp1), and double copy replacement strain (2X Dbp1) as determined by western blotting. One representative blot is shown. Data from four biological replicates is shown below; statistical significance was determined by an ordinary one-way ANOVA and corrected for multiple comparisons using Dunnett’s multiple comparison test with a Padj < .05 = *. (E) mRNA, FP, and TE measurements for *DBP1* and *DED1* coding sequences expressed in the 1x Dbp1, Ded1, and 2X Dbp1 strains. Two biological replicates are presented and statistical significance was determined by an ordinary one way ANOVA corrected for multiple comparisons with Dunnett’s multiple comparison test and a Padj < .05 = *, Padj < .01 = **, Padj < .001 = ***, Padj < .0001 = ****. (F) Positional data for ribosome footprints (FP) and mRNA over the *DHH1* locus. Data are shown for mitotic cells expressing either Ded1 (green) or matched levels of Dbp1 (blue). Ribosome profiling data are above, mRNA-seq below. Note increased ribosome footprints within the 5’ leader in Dbp1-expressing cells. (G) Metagene plots demonstrating ribosome occupancy over all genes in relation to start codons. FP reads were normalized to total mapped reads for each sample. A growth-defect-matched strain (ribosomal protein deletion *rpl26bΔ)* is shown for comparison to the slow-growing single copy Dbp1 replacement strain.

### Differential protein degradation or slow growth do not contribute to low Dbp1 abundance

The reduced Dbp1 synthesis levels measured by ribosome profiling, which resulted from reduced translation efficiency of the *DBP1* ORF compared to the *DED1* ORF, could alone produce the protein-level differences observed for the two helicases in our experiments (Figures 2H-J), but increased degradation of Dbp1 protein compared to Ded1 protein could also be a contributing factor. To test this, we first inhibited new protein synthesis, using treatment with cycloheximide, and measured Dbp1 or Ded1 protein levels by western blotting at intervals up to 3 hours. This experiment revealed no reduction in the protein abundance of either helicase (Figure 3B). Inhibition of proteasome-mediated degradation using treatment of mitotic cells expressing either Dbp1 or Ded1 with the drug MG132 also revealed no difference between the behavior of Dbp1 and Ded1 (Figure 3SB), suggesting that the difference in protein levels observed when the two proteins are expressed from *DED1’*s regulatory regions in mitotic cells is driven by differential translation of the ORFs encoding these two helicases rather than protein degradation.

The difference between Dbp1 and Ded1 protein synthesis was surprising, given that only the ORF differed between the two strains and most regulation of protein production is thought to be mediated by noncoding regions. While interesting, this difference also made it impossible to directly compare the activity of the two proteins *in vivo*. As this was our primary objective, we attempted to increase Dbp1 protein expression by modifying our original strategy (Figure 2A). In diploid cells, we inserted either one or two extra copies of the *DBP1* ORF flanked by *DED1* regulatory elements and inserted at the *TRP1* locus, in the Dbp1-expressing strains (“1.5X Dbp1” and “2X DBP1”, respectively; Figure 3C, 3SC). We found that this this strategy increased *DBP1* mRNA levels (Figure 3D, 3SD, S3E) roughly proportionally to the number of copies of the *DBP1* ORF present in the genome. When twice as many copies of *DBP1* ORF were supplied as *DED1* ORF, flanked by *DED1* regulatory regions in both cases, protein levels between the two helicases were indistinguishable by western blotting (Figure 3D, Figure 3SE). The higher Dbp1 protein levels also rescued mitotic growth to a degree that was similar to that seen in Ded1-expressing cells (Figures 3SG, 3SH).

The ratio of protein to mRNA was similar in diploid cells carrying either two, three, or four copies of *DBP1* (“1X”, “1.5X”, “2X” Dbp1, respectively) driven by *DED1* regulatory regions (Figure 3SB, 3SF), suggesting that protein degradation of Dbp1 was not altered by increased expression or by the increased cell growth that resulted from it. Moreover, cycloheximide treatment of 2X Dbp1, revealed no evidence for Dbp1 degradation, as seen for 1X Dbp1 cells (Figures 3B, Figure 3SI). These experiments suggested that increased Dbp1 protein observed in 2X Dbp1 cells was a result of increased mRNA production, rather than altered translation efficiency or degradation. We performed ribosome profiling in duplicate of diploid cell lines expressing either Ded1, 1X Dbp1, or 2X Dbp1 (Figure 3SC). This allowed us to confirm that total synthesis of Dbp1 was increased in 2X Dbp1 cells in correspondence with mRNA levels; thus the translational efficiency of the *DBP1* ORF was unchanged (Figure 3E). Because 2X Dbp1 cells grew comparably to Ded1-expressing cells (Figures 3SG, 3SH), this result also revealed that the poor translation of *DBP1* ORF was not a result of poor growth or decreased translation in 1X Dbp1 cells (Figure 3E, Figures 2B-E).

### Ded1 replacement with Dbp1 drives increased 5’ leader translation in mitosis

The accumulation of 5’ leader translation that is seen in cells lacking Ded1 (*ded1-ts;* (Guenther et al., 2018a)) was also seen, and to a similar degree, when low or Ded1-matched levels of Dbp1 were expressed, compared to cells expressing Ded1 (Figures 2F, 3F, 3G). This was apparent for a subset of individual highly expressed transcripts (Figure 3F) and if all data for annotated ORFs were analyzed in aggregate by metagene analysis (Figure 2F; 3G). The effect was moderate, but could not be explained by decreased bulk translation, as increased ribosome occupancy in 5’ leaders was not seen in cells lacking *RPL26B*, a condition that results in low bulk translation, equivalent to that seen in the “1x” Dbp1 cell line (Figure 3G; (Cheng et al., 2018a)). This result suggests that increased 5’ leader translation is inherent to Dbp1, compared to Ded1, and is not affected by bulk translation levels. It also suggests that Dbp1 is less effective at stimulating scanning and translation initiation than Ded1.

### The presence of Ded1 does not enhance Dbp1 translation

We considered that reduced translation of the *DBP1* ORF compared to the *DED1* ORF in exponentially growing mitotic cells could be based on the helicases themselves. Our experimental setup (Figure 2A, Figure 3C, Figure 3SC) resulted in a situation in which translation of the *DBP1* ORF could only be driven by Dbp1 and translation of the *DED1* ORF could only be driven by Ded1. If Ded1 is very efficient at supporting translation of its own transcript, this alone could explain the difference in translation that we observed for the two helicases. To directly test this possibility, we supplied diploid mitotic cells deleted for *DED1* at its endogenous locus with both Dbp1 and Ded1. In one strain, we inserted constructs encoding two copies of internally 3V5-tagged Dbp1 expressed from the *DED1* regulatory regions and one copy of untagged Ded1 expressed from its own regulatory regions at the *LEU2* and *TRP1* loci, respectively. In another strain, we inserted constructs encoding two copies of untagged Dbp1 expressed from the *DED1* regulatory regions and one construct driving internally 3V5-tagged Ded1 expressed from its own regulatory regions, again at the *LEU2* and *TRP1* loci (Figure 4A). We grew both strains mitotically and harvested cells to measure mRNA and protein during exponential growth phase. We performed qPCR to measure transcript abundance for the helicase tagged by 3V5 in each case, and anti-3V5 western blotting in parallel. To our surprise, Dbp1 protein, even when expressed from two genomic loci, was now lower than Ded1 protein expressed from one genomic locus (Figure 4A, 4SA). This is in contrast to the results seen when each helicase was expressed in isolation (Figure 3D, 3SC) and suggests that the presence of Dbp1 may increase scanning efficiency of the *DED1* 5’ leader or translation initiation of the *DED1* ORF; or the presence of Ded1 decreases translation initiation of the *DBP1* ORF. We concluded that the identity of the helicase present cannot explain the inherently poor translation of *DBP1* relative to *DED1* in mitotic cells.

**Figure 4:**
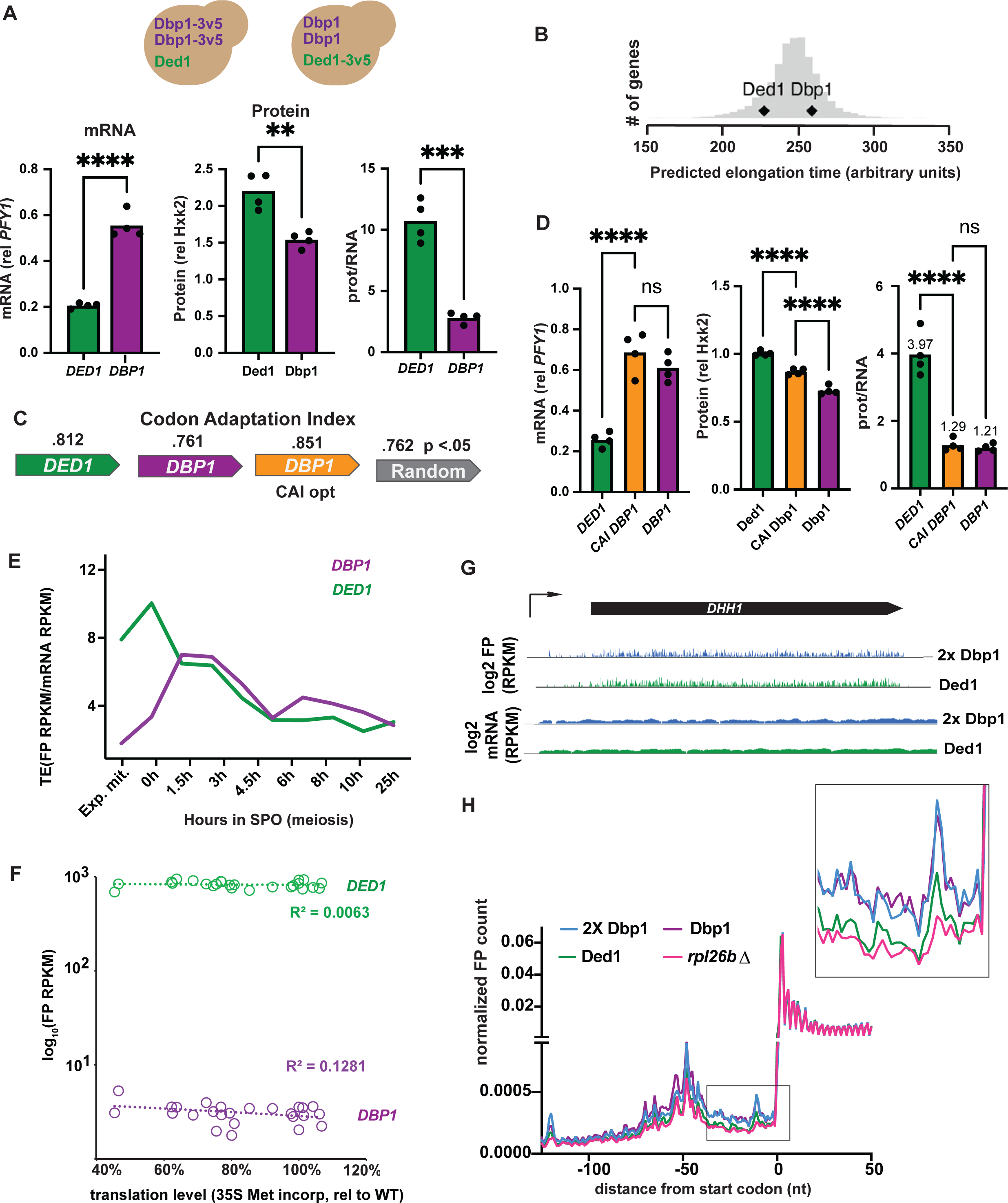
DBP1 codon optimization does not rescue its translation deficiency. (A) Schematic of strains expressing both Dbp1 and Ded1 helicase and the calculated pseudo TE for transcripts expressing *DBP1* or *DED1* in these strains. Levels of *DBP1* or *DED1* mRNA and protein as determined by RT-qPCR and western blot used for calculation of pseudo TE shown in Figure 3C. Data from four biological replicates is presented and statistical significance was determined by an unpaired t-test resulting in a two-tailed p < .05 = *, p < .01 = **, p < .001 = ***, p < .0001 = ****. (B) Predicted elongation time for DED1 and DBP1 coding sequences as predicted by a neural network model of translation speed. Predicted elongation times of all other endogenous yeast genes plotted in grey as a histogram (Tunney et al., 2018). (C) Codon Adaptation Index (CAI) for *DED1* and *DBP1* coding sequence as well as an amino acid composition matched set of random coding sequences as determined by the eCAI server (Puigbò et al., 2008). *DBP1* was codon optimized by replacing conserved amino acid positions with the codon from *DED1* if that codon was predicted to be better by CAI (codon optimized *DBP1* shown in orange). (D) Protein levels as determined by western blot, RNA levels as determined by RT-qPCR, and calculated pseudo TE of the *DBP1*, *DED1*, and codon optimized (CAI *DBP1*) *DBP1* coding sequences from (C). Data represent four biological replicates and statistical significance was determined by an ordinary one-way ANOVA corrected for multiple comparisons with Dunnett’s multiple comparisons test where a Padj < .05 = *, Padj < .01 = **, Padj < .001 = ***, Padj < .0001 = ****. (E) Translation efficiency (TE; ribosome footprint RPKM/mRNA RPKM) for *DBP1* and *DED1* during mitotic exponential growth and at time points through a meiotic time course (Cheng et al., 2018b) (F) translation (footprint RPKM) values for *DED1* and *DBP1* mRNAs relative to cellular translation level, as assessed by radioactive amino acid incorporation (Cheng et al., 2018a)

### Poor DBP1 ORF translation is independent of codon optimality

Ribosome profiling on either Ded1-, 1X Dbp1-, or 2X Dbp1-expressing diploid cells (Figures 3C, 3G; 3SD) revealed higher footprint density over the *DBP1* ORF (in the “2x Dbp1” strain) than the *DED1* ORF (in the “Ded1” strain), yet protein levels were comparable for the two helicases in these two cases. As we excluded protein degradation as a factor (Figures 3B, 3SB, 3SI), this suggested that slower translation elongation over the *DBP1* ORF than the *DED1* ORF could contribute to its relatively poorer corresponding protein expression level. A model to predict the translation elongation speeds for the *DBP1* and *DED1* ORFs based on codon use in exponentially growing cells determined that the codon makeup of *DBP1* ORF places it in the slowest 50% of genes while *DED1* ORF fell in the fastest 25% (Figure 4B;(Tunney et al., 2018)). Codon adaptation index (CAI) analysis, which scores codon optimality based on the overall codon composition of an ORF, also suggested *DBP1* to be sub-optimally coded relative to *DED1*, with its CAI value almost identical to a random sequence with similar GC/AT ratio and amino acid content (Figure 4C; (Puigbò et al., 2008; Sharp and Li, 1986)). To test whether increasing the codon optimality of *DBP1* could improve the amount of protein produced per mRNA, we recoded *DBP1* to surpass the CAI value of *DED1*. In homozygous diploid cells housing either *DED1, DBP1,* or the CAI-optimized *DBP1*, all flanked by *DED1*’s regulatory regions, we observed a partial increase in Dbp1 protein levels conferred by the increased CAI value of its ORF, but levels were not rescued to the level of Ded1 (Figure 4D). Moreover, there was no significant increase in the amount of protein produced per mRNA molecule as a result of increased *DBP1* CAI (Figure 4D, third graph), arguing that codon optimality is not the primary basis for poor translation of the *DBP1* mRNA, relative to that of *DED1*.

### Differential translation efficiency of DBP1 and DED1 is condition-specific

Relatively poor efficiency of translation seen for the *DBP1* ORF in our experiments could result from the artificial mRNA construct produced by fusing the *DBP1* ORF with *DED1* UTRs. This could either disrupt normal mRNA structure or create a new, inhibitory secondary structure. Examination of TE values in wild-type cells, however, did not support this model. Although *DBP1* mRNA is expressed at extremely low levels in exponentially growing mitotic cells, it is detectable by our mRNA sequencing depth, and a very low level of translation can also be observed by ribosome profiling under these conditions, in which wild-type transcripts are present for both *DED1* and *DBP1*. This results in consistently lower TE for *DBP1* relative to *DED1* in several mitotic datasets from our lab (Brar et al., 2012; Cheng et al., 2018b, 2018a), suggesting that poor mitotic translation is a natural property of the *DBP1* ORF.

The analysis of TE measurements from our previous datasets also revealed interesting condition-specificity to the TE of both *DBP1* and *DED1.* During meiosis, the TE for *DBP1* increases, whereas that of *DED1* decreases. In mid-meiosis, the TE of the two is almost identical, suggesting that some aspect of cellular environment enhances translation of *DBP1* mRNA and reduces translation of *DED1* mRNA. It was previously proposed that TE values can be affected by bulk translation levels within cells (Khajuria et al., 2018; Lodish, 1974; Mills and Green, 2017). Consistently, our study of ribosomal protein gene mutants showed that in exponentially growing mitotic cells, reduced bulk translation resulted in higher translation for transcripts with low TE in wild-type cells, and lower translation for transcripts with high TE in wild-type cells (Cheng et al., 2018a). Meiotic yeast cells show reduced bulk translation relative to mitotic cells (Brar et al., 2012). This, however, does not lead to the increase in *DBP1* or decrease in *DED1* TE seen in meiosis. The “0 hour” timepoint, with cells exposed to nutrient-poor media and primed to enter meiosis, exhibits much lower bulk translation than mid-meiotic timepoints (Brar et al., 2012) and yet the difference between *DED1* and *DBP1* TE is as high as in exponentially growing mitotic cells at this time, which have an even higher level of bulk translation (Figure 4E; (Brar et al., 2012; Cheng et al., 2018b)). Furthermore, we saw no correlation between translation levels of *DBP1* or *DED1* and bulk translation level across a spectrum of ribosomal protein gene mutations in exponentially growing mitotic cells, as assessed by ^35^S amino acid incorporation (Figure 4F;(Cheng et al., 2018a)). For this experiment, mRNA-seq RPKM were too low for *DBP1* for some samples to compare TE values for all, but values for those that could be calculated did not show changes. We concluded that cellular factors beyond bulk translation level changes cause the shifts in *DBP1* and *DED1* TE seen during meiosis.

### Loss of Dbp1 expression in meiosis leads to upregulation of Ded1 expression

Given the enhanced expression of Dbp1 in meiosis (Figure 1C), we examined the effects of Dbp1 loss in this context. Several studies have examined cells deleted for the *DBP1* ORF (Banroques et al., 2011, 2010, 2008; Berthelot et al., 2004; Sen et al., 2019), but all used a cassette insertion strategy that we recently reported to drive a strong and unexpected off-target effect via mis-regulation of mitochondrial translation that impacts cytosolic translation (Powers et al., 2022). We used a markerless CRISPR-based deletion strategy to create a *dbp1Δ* cell line that does not lead to this off-target effect (Powers et al., 2022) and displays no mitotic growth or translation defect, consistent with the lack of Dbp1 expression under these conditions. Surprisingly, we also observed little to no defect in meiotic progression (Figure 4SC) or meiotic translation (Figures 4SD, 4SE) in cells lacking Dbp1. Consistently, mRNA sequencing and ribosome profiling comparing WT and *dbp1Δ* cells in duplicate at early (3h) and mid-meiotic (6h) timepoints revealed few changes in mRNA, translation, or translation efficiencies genome-wide (Figures 4SF-H). These results were unexpected, given the apparent sensitivity of mitotic cell growth to helicase level (Figures 2SB, 2SC, 2SD; 3SG, 3SH) and the large contribution of Dbp1 to the Ded1/Dbp1 pool in meiosis (Figures 1B-1D). Examination of *DED1* expression provided an explanation: *dbp1Δ* cells showed a significant increase in translation of Ded1 (Figure 4SI, 4SJ), suggesting that meiotic cells compensate for low Dbp1 levels by upregulating translation of Ded1.

### Ded1 is more efficient at associating with ribosomes than Dbp1

The result that Dbp1 expression phenocopies the accumulation of 5’ leader ribosome occupancy observed with Ded1 deficiency (Figure 3G, (Guenther et al., 2018a)) suggests that Dbp1 is less active at enhancing translation than Ded1. This could be a result of poorer association with scanning ribosome subunits, presumably through weaker interactions with initiation factor partners (eIF4A, eIF4E, and/or eIF4G), poorer inherent helicase activity, or both. To test the first possibility, we performed mass spectrometry of ribosomes isolated from mitotic cells and cells at the meiotic timepoints shown in figures 1A and 4E (Figure 5A). We analyzed the translating ribosome fractions together, rather than 40S species, because most translation initiation occurs on mRNAs that are already being translated. We compared TMT10 (Tandem Mass Tag) mass spectrometry values for ribosome-associated fractions to those for matched total extract (Figure 5A). For general translation factors (such as eIF4A), the greater protein in extract in mitotic cells corresponded generally with greater protein associated with mitotic ribosomes. This was also true for Ded1 (Figure 5A). In the case of Dbp1, much higher total protein and ribosome associated protein was seen, as expected, in meiotic cells than mitotic cells. However, even at timepoints for which Ded1 and Dbp1 protein levels were equivalent in total extract, Dbp1 associated with ribosomes at substantially lower levels than Ded1 (Figure 5A).

**Figure 5:**
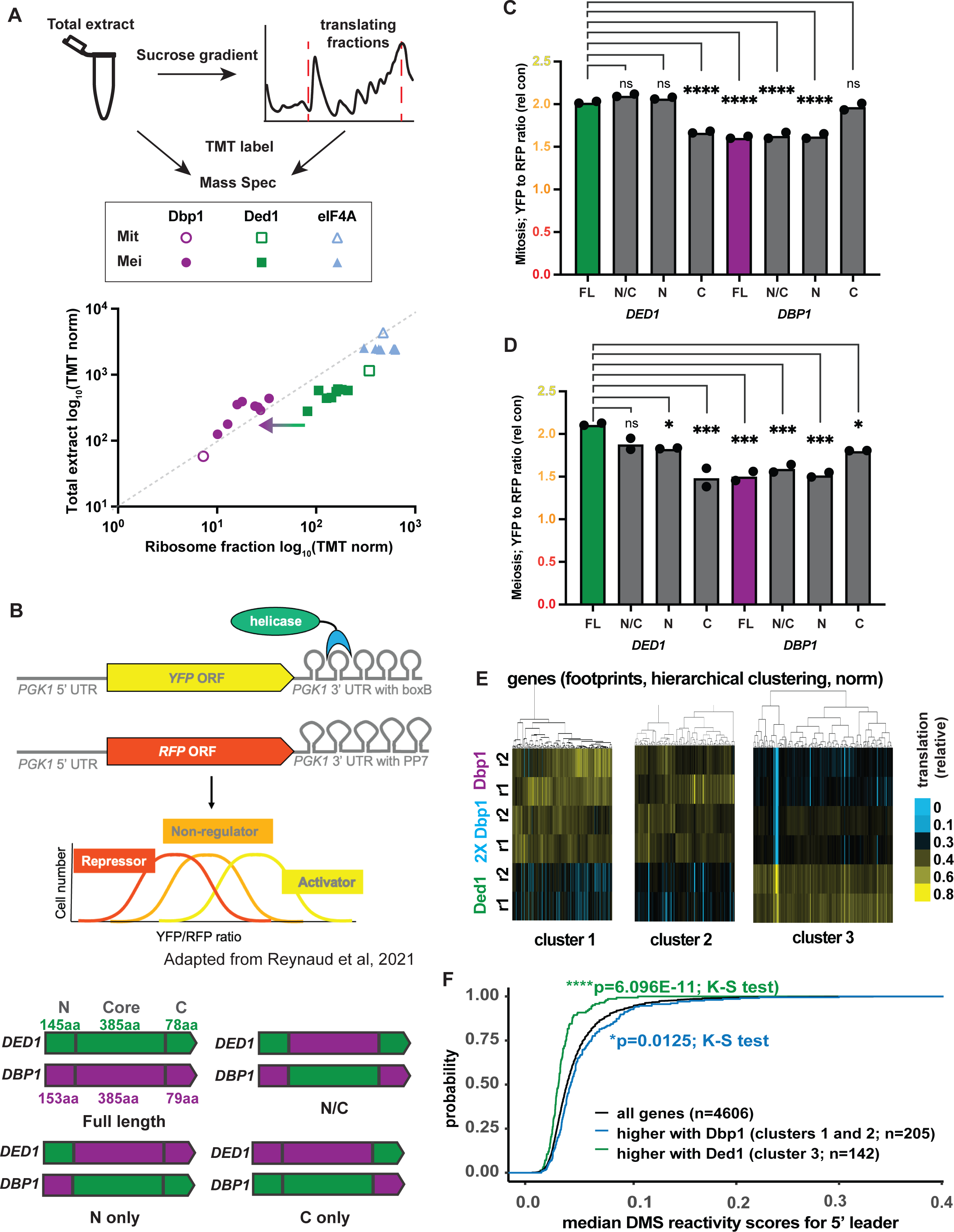
Dbp1 is less effective at driving translation activation than Ded1. (A) Levels of Dbp1, Ded1, and eIF4A RNA helicase that sediment with translating ribosome pools during mitosis or meiosis as determined by TMT-labeled mass spectrometry. Data represent the average of two biological replicates matched to FP data from (A) and total extract measurements matched to Figure 1A ribosome profiling from (Cheng et al., 2018b). Arrow indicates difference to note between Dbp1 and Ded1 ribosome association (B) Experimental set up for mRNA tethering assay. Figure is adapted from Reynaud et al., 2023. Schematics of full-length and chimeric proteins are shown below. (C,D) Ded1 or Dbp1 helicase was tethered to a YFP mRNA as outlined in (B). Mitotic (C) or meiotic (D) cells were analyzed by flow cytometry to determine their YFP to RFP ratio relative to a negative control. Data from two replicates are shown. Statistical significance determined by one-way anova corrected for multiple comparisons using Dunnett’s multiple comparison test where a Padj < .05 = *, Padj < .01 = **, Padj < .001 = ***, Padj < .0001 = ****. (E) Ribosome profiling data from the experiment schematicized in Figure 3C and 3SC were clustered for all transcripts measured (n=6219). The three clusters shown were identified as showing helicase-dependent differences and are shown here in isolation, with blue representing low translation and yellow representing high. Two replicates each are shown for Ded1-expressing mitotic cells, cells expressing matched levels of Dbp1 (2x) and cells expressing lower levels of Dbp1 (1x). (F) The median 5’ leader DMS reactivity scores from (Zubradt et al., 2017b) were determined for each of the transcripts in (E). Significance was assessed by K-S test, with those in clusters 1 and 2 combined for this analysis.

### Tethering experiments reveal reduced translation activation capacity for Dbp1

We next examined the ability of Dbp1 to enhance translation, using an mRNA tethering assay (Coller and Wickens, 2007; Reynaud et al., 2023). In short, we fused either Dbp1, Ded1, or a control Halo tag to the λN coat protein in cells expressing both a YFP reporter mRNA containing five boxB hairpins in its 3’ UTR (Figure 5B). An RFP reporter housing five PP7 hairpins in its 3’ UTR was also expressed in cells as a control and YFP to RFP ratio was assessed by flow cytometry. Tethering of the Ded1 helicase to the reporter mRNA increased the YFP/RFP ratio in mitotic cells, indicating that it behaves like a transcriptional activator (Figure 5B, 5C). This is consistent with published data, which revealed an N-terminal fragment of Ded1 to be among the most translationally activating protein regions in yeast (Reynaud et al., 2023). Dbp1 stimulated expression of the YFP reporter as compared to the negative control, but the magnitude of activation was lower than that seen with Ded1 (Figure 5C). Chimeric Dbp1/Ded1 helicases were also analyzed, by swapping the N-terminal, core, and C-terminal regions of the two helicases (Banroques et al., 2011). The N-terminus of Ded1, containing its eIF4E and eIF4A binding sites, was seen to be key to maximum translation activation by this assay (Figures 5B, 5C).

An advantage of the in-cell tethering approach is the ability to assay translational regulation under different in vivo conditions. Because Dbp1 is upregulated and Ded1 is downregulated during meiosis, we wondered whether Dbp1 may be more effective at activating translation in this cellular context. This was not the case. Meiotic cells displayed highly similar translational activation profiles to mitotic cells, with Ded1 leading to enhanced translation activation compared to Dbp1, and Ded1’s N-terminus playing a particularly important role in translation activation (Figures 5B, 5D). In meiotic cells, as opposed to mitotic, Ded1’s N-terminus alone could not fully support translational activation to the level seen with full-length Ded1, and a contribution of Ded1’s C-terminus alone was also seen in this cellular context (Figures 5B, 5D). Western blotting-based quantification of all helicases assayed revealed no relationship between total protein level and degree of translational activation, suggesting that the helicases were in excess of reporter mRNA (Figure S5A, S5B). In conclusion, these experiments reveal that Dbp1 is inherently less effective at stimulating translation than Ded1, even when artificially recruited to its mRNA target, and that this difference is driven by its N- and C- termini sequences.

### mRNAs with structured 5’ leaders are more efficiently translated by Ded1 than Dbp1

Ded1 function is important for stimulating translation generally, but multiple studies have also shown that its loss impacts a subset of mRNAs especially greatly. This subset is predicted to be more structured and contain longer 5’ leader regions than the average transcript (Guenther et al., 2018a; Gupta et al., 2018; Iserman et al., 2020; Sen et al., 2021, 2019, 2015). Our data that Dbp1 shows poorer association with ribosomes than Ded1 (Figure 5A), is less effective at stimulating translation (Figures 5B-5D), and that its expression in mitotic cells in place of Ded1 leads to enhanced translation within 5’ leaders (Figure 3G), all suggest that cells with mitotic replacement of Ded1 with Dbp1 behave as though they express a partial loss of function allele of *DED1*. If this is the case, we would expect mitotic cells expressing a Ded1-matched amount of Dbp1 to show reduced translation of mRNAs with structured 5’ leaders, compared to Ded1-expressing cells. To assess this, we examined the mRNA-seq and ribosome profiling datasets that we collected, comparing cells expressing either Ded1, low levels of Dbp1 (“1x”) or levels of Dbp1 that were matched to Ded1 (“2x”) (Figure 3G). The data were reproducible, as judged by replicate comparison (Figure S5C), but very few differences in mRNA or translation levels were observed when comparing 2x Dbp1 cells to Ded1-expressing cells (Figure S5D-F). The few changes that were seen did not correspond to those that are indicative of low overall translation levels (Figure S5G; (Cheng et al., 2018a)), suggesting that they were instead a result of differences in specificity of Ded1 versus Dbp1 in promoting translation initiation.

Using hierarchical clustering of all data together, we identified three clusters of interest. Two clusters (1 and 2, Figure 5E) contained mRNAs that were translated at a higher level when Dbp1 was expressed in place of Ded1. One cluster (3, Figure 5E) contained mRNAs that were translated at a higher level when Ded1 was expressed rather than Dbp1. The degree of difference in translation profiles between the two helicases in all three clusters was mild, with the 249 mRNAs in the two Dbp1-enhanced clusters showing only a 34% increase on average of ribosome footprints in 2x Dbp1 cells relative to those expressing Ded1 (excluding *DBP1* itself). Similarly, the 158 mRNAs in the Ded-enhanced cluster showed only a 20% average increase in translation in cells expressing Ded1 compared to 2x Dbp1-expressing cells (excluding *DED1* itself).

The mRNAs that showed enhanced translation when Ded1 was expressed, relative to Dbp1, were highly enriched for structured 5’ leaders, as assessed by in vivo mRNA structure determination by DMS-MaP-seq (Figure 5F; (Zubradt et al., 2017a)). This is consistent with reports that Ded1 is particularly important for translation of 5’ leaders with predicted structure (Guenther et al., 2018b; Gupta et al., 2018; Iserman et al., 2020; Sen et al., 2019) and our data that Dbp1 is less effective at driving translation initiation than Ded1 (Figure 5B-D). Interestingly, the mRNAs that showed enhanced translation in Dbp1-expressing mitotic cells, compared to Ded1-expressing mitotic cells, were significantly enriched for less structured 5’ leaders than a typical mRNA (Figure 5F), suggesting that some feature of Dbp1 causes it to be better than Ded1 at translating this small subset of mRNAs.

### Dbp1 cannot support robust cell growth at cold temperatures

Despite defects in translation of highly structured mRNAs, mitotic cells expressing Dbp1 at levels matched to Ded1 in wild-type cells display no measurable growth defect at standard laboratory growth temperatures (∼30°C; Figure S3H). We hypothesized that a growth defect may be unmasked by growing cells at cold temperatures, a condition in which mRNA structures would be expected to be stabilized, thus making Ded1 function of greater importance to cellular fitness. Indeed, we found that, while cells carrying four copies of *DBP1* (“2X”; Figure 3C) driven by *DED1* regulatory regions (Figure 3C) grew as well as those expressing Ded1 at 30°C, Dbp1 expression conferred a strong growth defect at 18°C (Figure 6A). This defect was more severe in diploid cells housing only two copies of *DBP1* (“1X”; Figure 3C) but this was not merely a result of slower growth in these cells, as cells lacking *RPL26B* displayed a similar mitotic growth defect to *1X DBP1* cells at 30°C, but a much less severe growth defect at 18°C (Figure 6B). The growth defect caused by Dbp1 expression during mitotic growth at 18°C was seen whether the helicase was internally 3V5-tagged or untagged (Figures 6A, 6C). C-terminal 3V5 tags, however, resulted in reduced growth of cells expressing either Ded1 or Dbp1 at 18°C, compared to untagged controls (Figure 6C). This is significant, given that most studies examining Ded1 function have relied on a C-terminally tagged version of the helicase. Ded1’s eIF4G binding site is at its extreme C-terminus (Aryanpur et al., 2019; Gao et al., 2016; Gulay et al., 2020; Hilliker et al., 2011; Senissar et al., 2014), which could be occluded by use of a C-terminal tag, and could thus explain this result. For this reason, we used internal tagging of these helicases for all experiments for which it was possible (excluding the tether experiments, as noted; Figure 5B-D).

**Figure 6:**
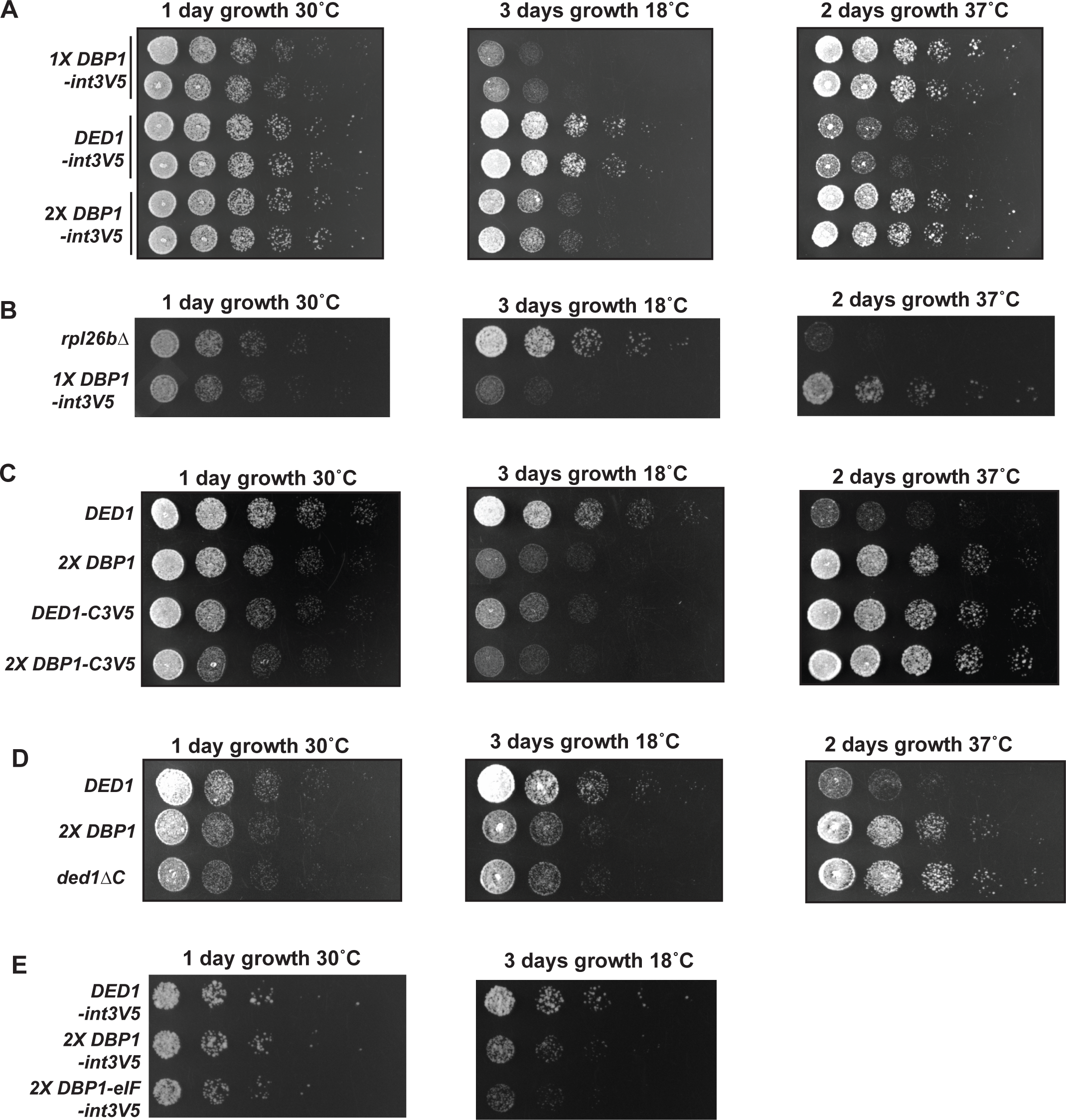
Dbp1 fails to support growth at low temperature, growth quelling at high temperature. (A-D) Growth of diploid cells on rich media (YEPD) plates is shown, using serial 1:5 dilution. Plates were grown at either 30°C, 18°C, or 37°C. (A) Cells compared include internally 3V5-tagged helicase, with either Ded1, matched Dbp1 (2x), or low level (Dbp1) cell lines shown. (B) Growth matched cells deleted for *RPL26B* or with the *DED1* ORF replaced homozygously (“*1X DBP1*”). (C) Untagged strains expressing either matched levels of Ded1 or Dbp1 are shown above. Below are similar strains carrying a C-terminal 3V5 tag. (D) Untagged strains expressing either matched levels of Ded1 or Dbp1 are shown above. Below are cells expressing Ded1 lacking its 14 C-terminal amino acids. (E) Cells were grown as in (A-D) but these are haploid rather than diploid cells. Shown are strains expressing either matched levels of Ded1, Dbp1, or a chimeric Dbp1 lacking six regions characterized to confer eIF binding to Ded1 (Gulay et al., 2020).

The C-terminus of Ded1 is required for its robust interaction with eIF4G (Gulay et al., 2020). In light of this, and the apparent defects in Ded1 function seen with C-terminally tagged protein (Figure 6C), we evaluated the effects of truncations within this region. Although previous studies showed that haploid cells expressing a tagged version of Ded1 lacking its C-terminal 43 amino acids did not alone display a growth defect at 16°C or 18°C (Gulay et al., 2020; Hilliker et al., 2011), we found that in diploid cells expressing internally 3V5-tagged Ded1, loss of the 43 amino acids of Ded1 could not support viability, even at 30°C. An internally tagged version of Ded1 lacking only its 14 C-terminal amino acids, which removed most of the eIF4G interaction region (Gulay et al., 2020), was viable at 30°C, but cells expressing it displayed a growth defect relative to full-length protein that was exacerbated when cells were grown at 18°C (Figure 6D). This result is consistent with the model that Ded1’s C-terminus is important for supporting its role in activating translation.

The helicase tethering experiments that we performed to assess translation-activating capacity of Dbp1, Ded1, and chimeric helicases suggested that the N- and C-termini of Ded1 were particularly important for activating translation (Figure 5B-D). The poor growth seen in Dbp1-expressing cells at 18°C, which was partly phenocopied by a version of Ded1 lacking its C-terminus (Figure 6C, 6D), the divergence of Dbp1 and Ded1’s N- and C-terminal sequences (Figure 1A), and the relatively poor recruitment of Dbp1 to ribosomes relative to Ded1 (Figure 5A), led us to wonder if Ded1’s superior ability to activate translation relative to Dbp1 is based on its eIF4F-interacting intervals. If so, supplying Dbp1 with the six peptide segments shown to be important for Ded1 binding to eIF4A, eIF4E, and eIF4G should improve Dbp1’s ability to support growth at 18°C (Gulay et al., 2020; Hilliker et al., 2011). We found that this was not the case (Figure 6E), suggesting that while these residues are needed for robust eIF4F component binding to Ded1 (Gulay et al., 2020), they may not be sufficient to supply full binding to Dbp1. Our analyses of cold temperature growth of mitotic cells expressing Ded1 or Dbp1 led us to conclude that expression of the Dbp1 helicase is unable to support growth as well as Ded1 under these conditions, presumably owing to its inferior ability to drive PIC scanning.

### Dbp1 cannot suppress cell growth at high temperatures

Ded1 is best known for its ability to activate translation, but it is also capable of driving translational repression. This can be seen when Ded1 is overexpressed, and is also associated with stress granule formation in response to external stressors—including glucose starvation or high temperature—with these granules leading to repression of translation of the majority of “housekeeping” mRNAs and reduced cell growth (Aryanpur et al., 2022; Hilliker et al., 2011; Iserman et al., 2020). Because the heat-induced condensation of Ded1 into stress granules is regulated by its disordered N- and C-termini, which differ from Dbp1, we investigated the ability of Dbp1 to support reduced growth at high temperatures. To test this, we grew diploid cells expressing either Dbp1 or Ded1, driven by *DED1*’s regulatory regions, on rich media (YEPD) plates at 37°C. We found that, in contrast to Ded1-expressing cells, cells expressing Dbp1 were unable to halt growth under these conditions, whether Dbp1 was untagged or internally tagged (Figure 6A, 6C). Ded1’s ability to drive condensation is enhanced by its C-terminus, and consistently, loss of only its 14 final amino acids phenocopies replacement of the entire protein with Dbp1 (Figure 6C, 6D; (Iserman et al., 2020)). These experiments revealed Dbp1 to be deficient in quelling growth under heat stress. We conclude that, although Dbp1 can largely perform similar functions in translation initiation as Ded1, as previously reported (Berthelot et al., 2004; Jamieson and Beggs, 1991; Sen et al., 2019), it is less effective in vivo at activating translation, and in halting translation in response to an external stressor.

## Discussion

Ded1 has been highly studied but its paralog, Dbp1, has not. The few in vivo studies of Dbp1, along with its sequence similarity to Ded1, have suggested that it supports translation initiation in manner that is similar to Ded1. These studies, however, have all been complicated by either a surprising off-target effect of selection-cassette-based replacement of the *DBP1* ORF or have used multicopy plasmid expression of Dbp1, leading to non-endogenous levels (Banroques et al., 2011, 2010, 2008; Berthelot et al., 2004; Powers et al., 2022; Sen et al., 2019). Low copy plasmid expression of Dbp1 was not able to substitute for Ded1 in supporting mitotic growth, whereas high copy expression was (Banroques et al., 2011; Jamieson and Beggs, 1991; Sen et al., 2019), and Ded1 switches from acting as a translational activator to act as a translational repressor when overexpressed. Together, these findings suggest that controlling for the levels of these DEAD-box helicases is important for elucidating their in vivo functions. Our study, which used single copy genomic replacement of *DBP1* and *DED1*, was the first to monitor this. Our findings show that expression levels matter for experimental interpretation, and that the cell controls them highly, in multiple ways. Through these protein-level-controlled experiments, we were able to test Dbp1 versus Ded1 functions in bulk translation, and to identify transcript-specific effects that were independent of expression level differences.

Most Ded1 is not ribosome-associated, making it somewhat surprising that the roughly 50% lower level of Dbp1 expression observed in mitotic cells in which the *DED1* ORF is replaced with the *DBP1* ORF leads to defective translation and cell growth. This could be explained by high free levels of the helicase serving an important role in driving sufficient Ded1 or Dbp1 to eIF4F-bound mRNAs and/or PICs but it also seems that Dbp1 is inherently less efficient at associating with ribosomes, at stimulating translation even when tethered to a reporter transcript, at activating translation of highly structured 5’ leaders, and at bypassing suboptimal translation initiation codons within 5’ leaders. All of these results argue that, even when comparing matched protein levels, Dbp1 is less effective supporting translation than Ded1, in a manner that depends on the divergent N- and C-termini of the two helicases (containing Ded1’s eIF4F factor binding sites; (Aryanpur et al., 2019; Gulay et al., 2020)). Most of the defects seen in mitotic cells expressing an equivalent level of Dbp1 in place of Ded1 are modest, suggesting that at least in rich exponential-phase mitotic growth, the two helicases are largely comparable in their translation-stimulating function. However, these functional differences may be of greater importance in suboptimal growth conditions. As one example, cells expressing *DBP1* in place of *DED1* at 18°C display a profound growth defect, potentially resulting from the more stable RNA secondary structures that exist at low temperatures.

One of the most surprising findings of our study was the poor translation of *DBP1*’s own mRNA compared to that of *DED1*, an effect dependent on the ORF sequences of the two transcripts. This difference in translatability remained even when *DBP1* codon optimality was modified to exceed *DED1*. It is condition-specific, with the translation efficiency of *DBP1* rising and *DED1* dropping during meiosis. One possibility is that the mRNA secondary structure of one or both ORFs may be a contributing factor; another is that a trans-factor drives the inherent differences between the ORFs in mitosis or their similarity in meiosis. Considering the relatively poor translation efficiency of Dbp1 in mitosis together with its relatively poor performance as an initiation helicase is interesting. During mitosis, even when the transcripts are present at high abundance, Dbp1 expression is severely dampened compared to Ded1. This enforces preferred PIC access to the more efficient initiation helicase, Ded1, in mitotic cells. Given that cellular growth rate in rich media is thought to be limited by translation rate, this should also enforce maximal cell growth. In contrast, in meiosis, translation levels are inherently lower and cells are also able to express Dbp1—the less efficient helicase at driving translation initiation—more efficiently.

Cells expressing *DBP1* in place of *DED1* display enhanced growth at 37°C. The cessation of mitotic cell growth at high temperatures has been shown to be, in part, based on changed behavior of Ded1, which forms stress granules that result in poor translation of housekeeping mRNAs and enhanced translation of transcripts that encode stress-response factors (Aryanpur et al., 2022; Hilliker et al., 2011; Hondele et al., 2019; Iserman et al., 2020). This allows cells to respond to the new and stressful condition of heat stress. The inability of cells expressing Dbp1 to halt growth under these conditions suggests that Dbp1 is less capable of translation inhibitory functions function than Ded1. Together, our results comparing the functions of the two helicases support the model that Ded1 is a “high performance” initiation helicase, perhaps akin to an F1 vehicle. It can support maximal translation and can allow cells to quickly halt translation, while Dbp1 is less powerful at both activating and braking functions, more like a family sedan.

If Dbp1 is less effective than Ded1 at stimulating translation and quelling growth in response to stress, what is its purpose in cells? The conditions in which it is expressed may provide insight into this question (Figure S1A). All reflect chronic stress conditions, in which cellular resources are limiting for long timespans that exceed typical mitotic doubling times (Gasch et al., 2000). In the case of meiosis, a cellular differentiation program that is stimulated in yeast by extreme nutrient deprivation, bulk translation levels are lower than during mitotic exponential growth (Brar et al., 2012), and thus a helicase that is only moderately efficient at stimulating translation may be tolerated, perhaps even preferable. It may also not be advantageous for cells to stop meiotic progression in response to an acute external stressor, as they cannot return to mitotic growth after a restriction point relatively early in the meiotic program. Halting translation would simply delay gamete formation without hope of improved conditions. Further experiments that are complementary to those presented in this manuscript will be important to identify specific conditions in which Dbp1 expression is preferable to Ded1.

The increase in 5’ leader translation seen in mitotic cells expressing Dbp1 in place of Ded1 phenocopies the increase in translation initiation at near-cognate codons in 5’ leaders seen in cells deficient for Ded1 function. Spurious translation initiation would be expected to generally decrease the overall efficiency of synthesis of proteins encoded downstream of these sites but it could also offer an advantage under certain circumstances. We recently showed that translation initiation within 5’ leaders is common in meiosis and can result in production of alternative N-terminally extended protein isoforms (Brar et al., 2012; Eisenberg et al., 2020). Such isoforms are known to be important, in some cases, for dually targeting a protein product to an additional subcellular location. The enhanced near-cognate initiation seen in meiosis is in part dependent on low eIF5A expression, but the low Ded1 and high Dbp1 expression under these conditions may also contribute. The set of factors that drive this increase in production of noncanonical protein isoforms diversifies the proteome in meiosis, and may support cellular functions that are important in this cellular context, and that enhance meiotic fidelity. It is possible that a transition from Ded1 to Dbp1 during meiosis allows cells to ramp up 5’ leader initiation while still allowing for enough translation to support progression through the meiotic program.

## Acknowledgements

We thank Paige Diamond for sharing global structure analysis file used for Figure 5F, Liana Lareau for elongation time plot for *DBP1* and *DED1* ORFs (Figure 4B), and BrÜn lab members and for helpful comments on the manuscript. This work was supported by National Institutes of Health funding to G.A.B. (R35GM134886) and M.J. (R35GM128802). E.N.P was funded by an NSF predoctoral Fellowship.

## Materials and Methods

### Strains

All strains used in this study were of derivatives of Saccharomyces cerevisiae of strain background SK1, outside of those used for tethering experiments which were of BY or hybrid SK1 and BY background. All genome edited transformants were backcrossed at least once prior to use. To construct the *ded1Δ* allele we first cloned a single copy of the *DED1* ORF C terminally tagged with 3v5 and ∼1kb of upstream and downstream regulatory sequence into a single integration vector and inserted it into the *LEU2* locus. This strain was then transformed with a plasmid encoding Cas9 and a sgRNA targeting the C terminus of *DED1* that is unable to target the C terminally tagged Ded1 allele inserted at the *LEU2* locus. The Cas9 editing plasmid was transformed alongside a linear repair template that deletes the *DED1* ORF. The *ded1Δ* was backcrossed and all subsequent rescue strains were created by transforming a wild-type strain with the *LEU2* integrated helicase, backcrossing, then crossing the rescue allele to the ded1Δ. Internal helicase tags were designed by finding sites that tolerated insertions within the Ded1 helicase by looking at sequence alignments to homologs from other species. Possible insertion sites were confirmed to be un-structured and surface exposed by examining their location on a published Ded1 structure. Tag functionality was determined by comparing the growth of tagged and untagged helicases under all growth conditions tested in this study. The *dbp1Δ* allele was created by transforming a wild-type haploid with a Cas9 editing plasmid expressing a sgRNA targeted to the *DBP1* ORF alongside a linear repair template that deletes the *DBP1* ORF. This haploid was backcrossed to wild-type yeast and subsequent haploids were mated to create a diploid.

### Mitotic growth conditions

For all mitotic growth experiments except for the tethering assays, yeast were grown in rich media (YPD 2% dextrose). Unless otherwise specified, cells were grown overnight at 30°C then back diluted to .05OD_600_/mL on the day of the experiment and grown until they reached exponential growth rates ∼.6OD_600_/mL. For meiotic experiments, cells were grown as previously described in Powers et al., 2021. For serial dilution experiments, cells were grown overnight or for the indicated length of time shaking at 30°C. Cells were diluted to .2OD_600_/mL then serially diluted 1:5 and 3uL of cells from each dilution were loaded onto plates.

### mRNA tethering assay

Each helicase of interest was C-terminally tagged with the lambda N RNA-binding domain, to tether it specifically to a YFP mRNA containing the boxB binding site (within the 3’ UTR) (Reynaud et al., 2023). Helicase fusions were also tagged with BFP and the 3xFLAG peptide sequence so their expression could be monitored. CEN plasmids encoding these constructs and marked with the S. pombe HIS5 allele for selection were transformed into a diploid strain of the BY background which contained heterozygous constructs expressing either YFP including the boxB sequence, or mCherry (with no boxB sequence) integrated at the *URA3* locus. Two independent transformants containing each helicase tether construct were then grown overnight, and then diluted to 0.1 OD_600_/mL the next day. Once the cells had grown to 0.6 OD_600_/mL, samples were fixed in 4% paraformaldehyde for 20 minutes room temperature for flow cytometry or incubated in 5% trichloroacetic acid overnight at 4°C for western blot analysis. All strains were grown in SC-His and at 30°C. For meiotic experiments, the BY strain expressing the YFP or mCherry constructs was backcrossed 3 times to wild-type yeast of the SK1 background. The resulting strain of mostly SK1 background had the same YFP and mCherry constructs as used in the vegetative experiments and was transformed with helicase expression constructs. Transformants were grown overnight in SC-His , diluted to 0.25 OD_600_/mL in BYTA and grown overnight again, then diluted into SPO media at 1.9 OD_600_/mL. After 4.5h in SPO, western blot samples were collected as described above to assess helicase abundance, and after 5h in SPO samples were collected for flow cytometry as described above.

### Western blotting

Samples were prepared by TCA precipitation and extraction as described in section 1.5.4 of this thesis with the following differences. Blots were incubated with a rabbit anti FLAG antibody (1:1000, RRID; AB_2217020, 2368, Cell Signaling Technology), rabbit anti-hexokinase antibody (1:10,000, Rockland 100-4159), mouse anti-V5 (1:2000, Invitrogen R960-25), rat anti-tubulin antibody (1:10,000, Serotec, RRID:AB_325005). Blots were blocked in PBS 5% NDFM.

### Ribosome profiling and polysome profiling

Ribosome profiling and polysome analysis was performed as in (Powers and Brar, 2021). Briefly, cells were treated with cycloheximide for 30s then filtered and flash frozen. Extracts were milled under cryogenic temperatures and stored at -80°C in aliquots. RNA extracted from monosomes was extracted and fragments ∼28-32nt were collected. Libraries were prepped using linker ligation and rRNA fragments were depleted from samples using biotinylated anti rRNA oligos. Samples were sequenced using 50nt single end reads on a HS4000 (Figures 3.3-3.4) or using 50nt single end reads on a NovaSeq 6000 (Figures 3.6-3.7). Matched mRNA-seq libraries were prepared with the same library prep protocol with the following changes. Poly A selection was used to isolate mRNA from extracted total RNA samples which were subsequently fragmented and libraries were created with fragments ∼35-80nt. No rRNA depletion was performed on these samples.

### mRNA-seq and ribosome profiling analysis

Adaptor sequences were trimmed off reads which were then aligned to the yeast genome as previously described in (Cheng et al., 2018b). Reads per ORF were counted and RPKMs were calculated for each gene by normalizing the raw reads to the sum of reads per sample and the gene length. Differential expression analysis was performed on raw read counts using DESeq2. Genes with a P_adjusted_ value of less than or equal to .05 were considered significantly different between samples. Sequencing data shown in plots represents the average of 2 or 3 biological replicates with all data shown. To do GO analysis, significantly regulated genes were split into those upregulated or downregulated in the Dbp1 expressing strains compared to their Ded1 expressing partner. The top 5 enriched GO terms for each category are reported.

### Analysis of meiotic progression

Cells were prepared to undergo meiosis as described in Powers et al., 2021. For analysis of meiotic divisions, cells were collected at the noted timepoints by overnight treatment with 3.7% formaldehyde at 4°C. Cells were resuspended in KPi buffer (100 mM potassium phosphate, pH 6.4) and adhered to poly-L-lysine treated glass slides and membranes were permeabilized by brief treatment of 70% ethanol on slide. Ethanol was aspirated and when wells were dry VectaShield Antifade Mounting Medium with DAPI (Vector Labs) was added. Slides were sealed with a coverslip and used to count nuclei.

### Mass spectrometry of mono/polysomes

Extract used for total protein quantification in (Cheng et al., 2018b) was subjected to sucrose gradient fractionation and approximately 6 ml of material was collected per sample, containing the monosome/80S and polysome fractions were collected. This was done on biological replicate samples for all 10 conditions analyzed. Proteins were processed by the FASP protocol (Wiśniewski et al., 2009). Briefly, 100 μl of sample was mixed with 400 μl 8M Guanidine Hydrochloride, then loaded onto a Nanosep Omega 10K column and spun at 14,000g till dry. Then the sample was washed twice with 400 μl Urea Buffer (8M Urea, 50mM Tris/HCl (pH8), 75mM NaCl, 1mM EDTA) and spun till dry. 100 μl Urea Buffer was added. Disulfide bonds were reduced with 5 mM dithiothreitol and cysteines were subsequently alkylated with 10 mM iodoacetamide. Afterwards the samples were spun till dry and 200 μl of 1:4 diluted (dilute with 50mM Tris/HCl (pH8)) urea buffer was added together with Trypsin and LysC at a ratio of 1:100 to total protein. The samples were then incubated at 25°C overnight. The column was transferred to a new 1.5 ml Eppendorf tube and the digested sample was collected by a spin at 14,000g. Tryptic peptides were desalted on C18 StageTips according to (Rappsilber et al., 2007) and evaporated to dryness in a vacuum concentrator. Desalted peptides were labeled with the TMT-10plex mass tag labeling reagent according to the manufacturer’s instructions (Thermo Scientific) with small modifications. Briefly, 0.2 units of TMT-10plex reagent was used per 10 μg of sample. Peptides were dissolved in 30 μl of 50 mM Hepes pH 8.5 solution and the TMT-11plex reagent was added in 12.3 μl of MeCN. After 1 h incubation the reaction was stopped with 2.5 μl 5% Hydroxylamine for 15 min at 25°C. Differentially labeled peptides were mixed for each replicate (see mixing scheme below) and subsequently desalted on C18 StageTips (Rappsilber et al., 2007), evaporated to dryness in a vacuum concentrator and reconstituted in 15 μl of 3% acetonitrile and 0.1% formic acid.

| **TMT mix** | **Sample** | **TMT label** |
| --- | --- | --- |
| 1 | *vegetative* | 126C |
| 1 | *Meiosis 0h* | 127N |
| 1 | *Meiosis 1.5h* | 127C |
| 1 | *Meiosis 3h* | 128N |
| 1 | *Meiosis 4.5h* | 128C |
| 1 | *Meiosis 6h* | 129N |
| 1 | *Meiosis 8h* | 129C |
| 1 | *Meiosis 10h* | 130N |
| 1 | *Spores 25h* | 130C |
| 1 | *MATa/a 4.5h in sporulation medium* | 131N |
| 2 | *vegetative* | 131N |
| 2 | *Meiosis 0h* | 130C |
| 2 | *Meiosis 1.5h* | 130N |
| 2 | *Meiosis 3h* | 129C |
| 2 | *Meiosis 4.5h* | 129N |
| 2 | *Meiosis 6h* | 128C |
| 2 | *Meiosis 8h* | 128N |
| 2 | *Meiosis 10h* | 127C |
| 2 | *Spores 25h* | 127N |
| 2 | *MATa/a 4.5h in sporulation medium* | 126C |

LC-MS/MS analysis on a Q-Exactive HF was performed as previously described (Cheng et al., 2018c; Keshishian et al., 2015). Briefly, around 1 μg of total peptides were analyzed on an EASY-nLC 1000 UHPLC system (Thermo Fisher Scientific) coupled via a 20 cm C18 column ID picofrit column (New Objective, Woburn, MA) packed in house with Reprosil-Pur C18 AQ 1.9 μm beads (Dr. Maisch, GmbH, Entringen, Germany) to a benchtop Orbitrap Q Exactive HF Plus mass spectrometer (Thermo Fisher Scientific).

### Quantification and Statistical Analyses for Mass Spectrometry

All mass spectra were analyzed with the Spectrum Mill software package v4.0 beta (Agilent Technologies) according to (Mertins et al., 2013) using the yeast Uniprot database (UniProt.Yeast.completeIsoforms.UP000002311.20151220; strain ATCC 204508 / S288c). For identification, we applied a maximum FDR of 1% separately on the protein and peptide level and proteins were grouped in subgroup specific manner. We required at least 1 spectral count from a unique peptide for protein identification and for protein quantification per replicate measurement. Note that the S288C UniProt dataset was used, because we are not aware of an equivalently complete protein dataset for SK1, and due to poorer sequencing depth and annotation of this genome relative to the reference, our attempt to create one excluded many proteins. This presumably caused us to miss capture of some proteins for which the quantifiable peptides are not identical in the two strains, but should not cause artifacts in our correlation measurements, because all measurements are relative among timepoints.

Finally, we normalized the Spectrum Mill generated intensities such that at each condition/time point the TMT intensity values added up to exactly 1,000,000, therefore each protein group value can be regarded as a normalized microshare (we did this separately for each replicate for all proteins that were present in that replicate TMT mix).

### RT-qPCR

RNA was isolated from the gradient fraction samples or the cell pellets by hot acid phenol extraction, DNase-treated with Turbo DNase (Invitrogen), and purified with phenol extraction. The RNA samples were adjusted to similar concentration and 150-250 ng of RNA was used in reverse transcription with Superscript III (ThermoFisher). Transcript levels were quantified on a StepOnePlus Real-Time PCR system using the SYBR green PCR mix (ThermoFisher), and normalized to the total RNA concentrations. CT values were first transformed to fold changes (according to the standard curve of the primer pair). Primers were tested for specificity and linear-range detection.

### Cycloheximide conditions

Cells grown in YEPD were treated with cycloheximide to a final concentration of 100ug/mL from a 500X stock in ethanol

### MG132 conditions

Cells grown in YEPD were treated with MG132 to a final concentration of 100uM from a 1000X stock in DMSO.

### Data accessibility

Processed and raw mRNA-seq and ribosome profiling data will be deposited at NCBI GEO and made publicly accessible at the time of publication.

