## Supplementary material for "Dbp1 is a low performance paralog of RNA helicase Ded1 that drives impaired translation and heat stress response": Figure 1S

A

| Condition | *DBP1* | *DED1* |
|---|---|---|
| Mitotic exponential | VERY LOW | HIGH |
| Meiosis (highly synch, rel to mitotic) | ⬆ 35.4X | ⬇ 5.8X |
| Stationary Phase (3-5 days) | ⬆ 5.3X | ⬇ 3.5X |
| Chronic Low Nitrogen (3 days) | ⬆ 3.0X | ⬇ 1.8X |
| Post-Diauxic Shift | ⬆ 1.8X | ⬇ 2.0X |

B

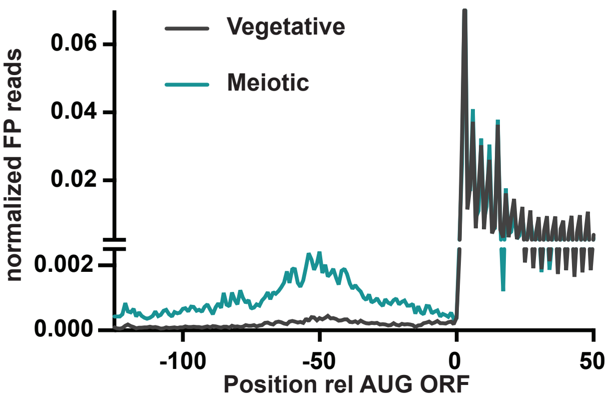
