## Supplementary figures and images for "Dbp1 is a low performance paralog of RNA helicase Ded1 that drives impaired translation and heat stress response"

### Figure 2S

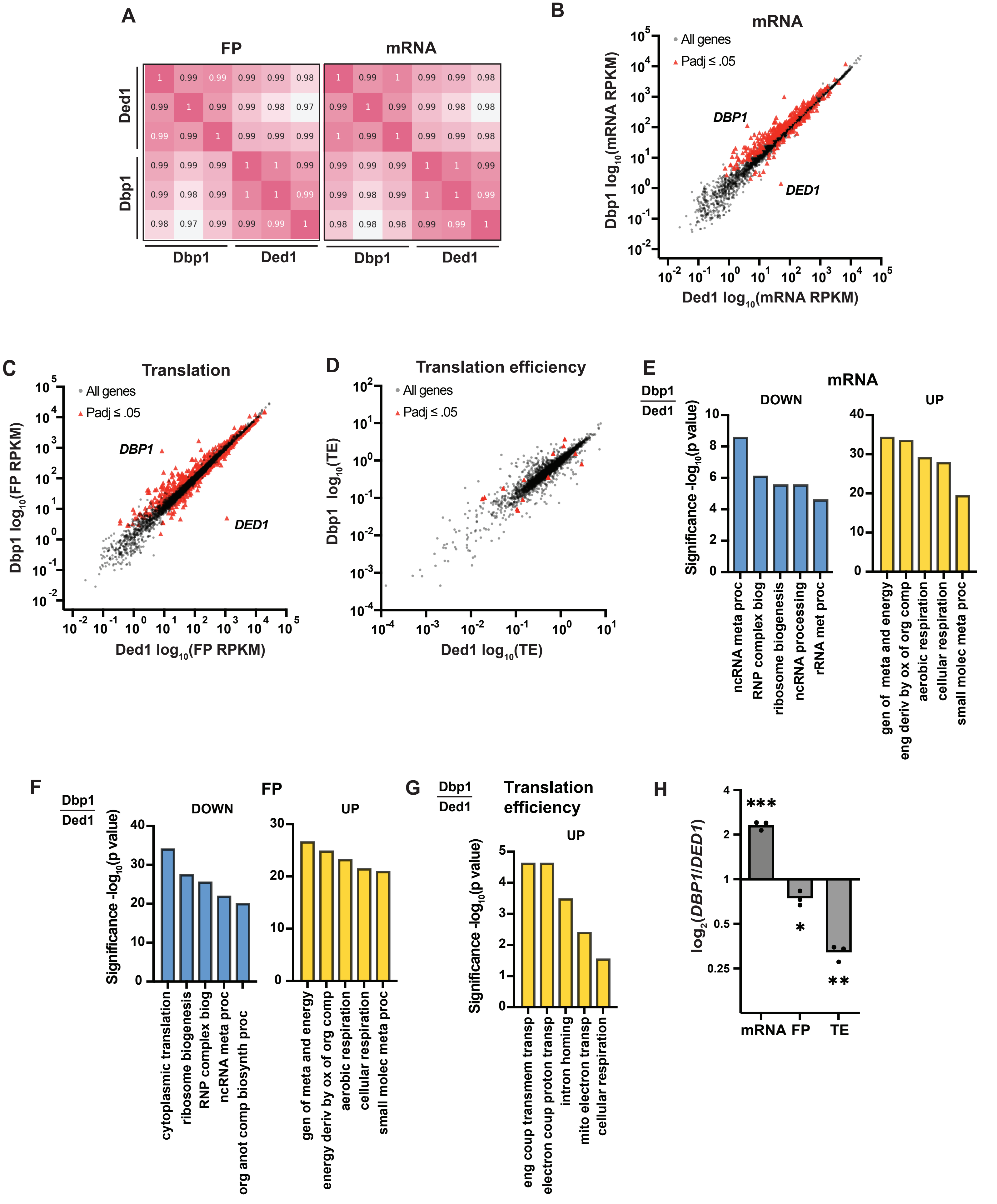

### Figure 3S

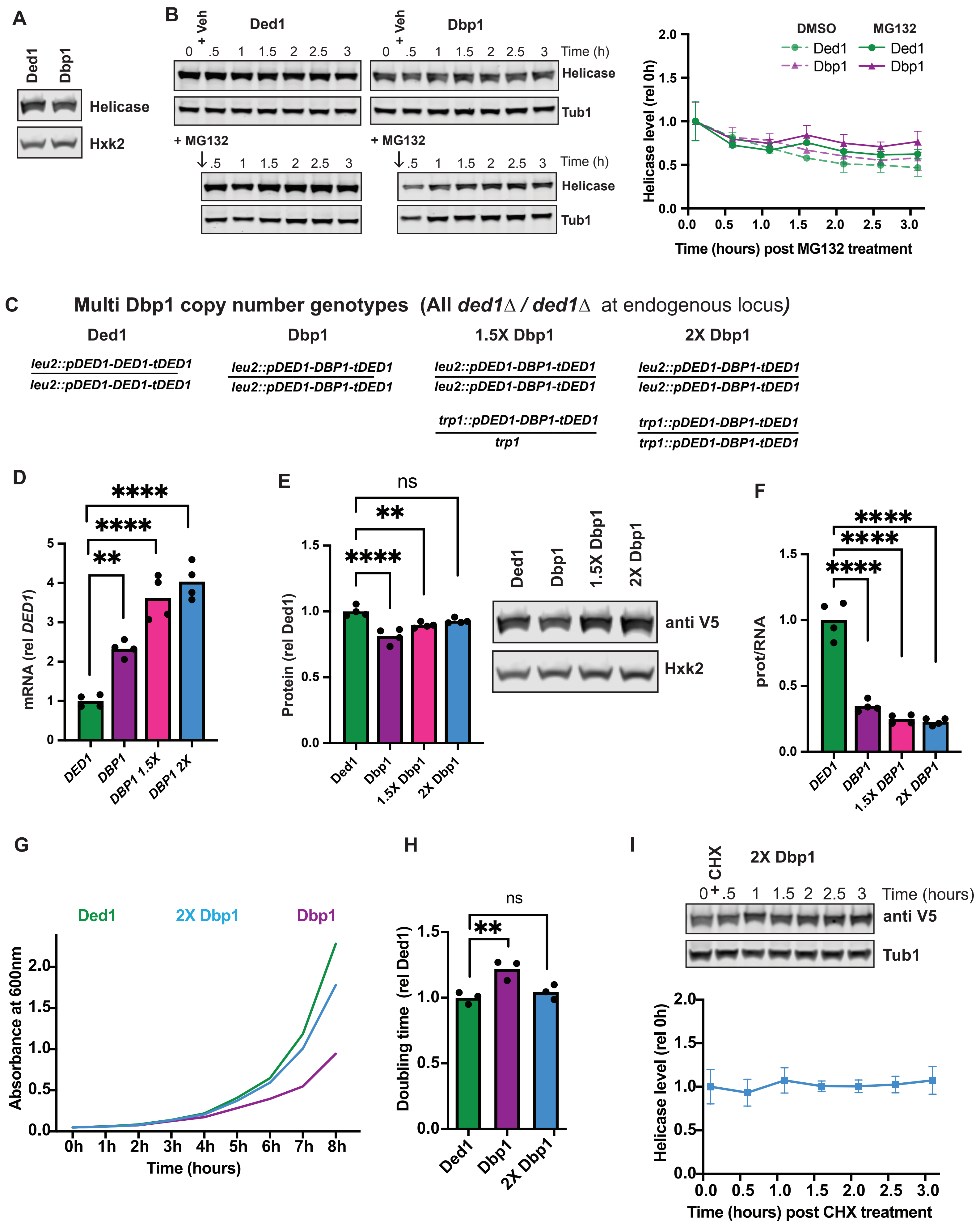

### Figure 4S

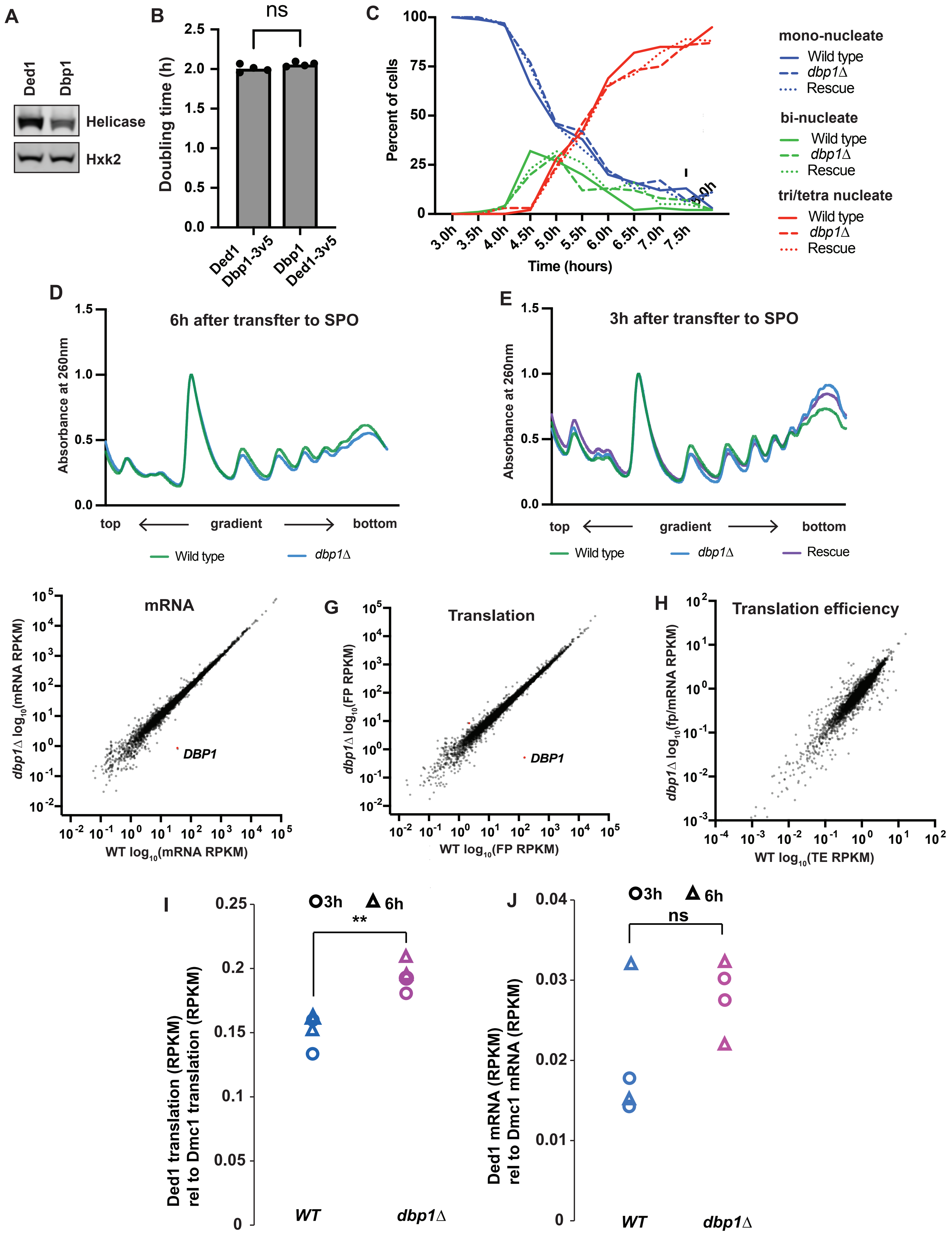

### Figure 5S

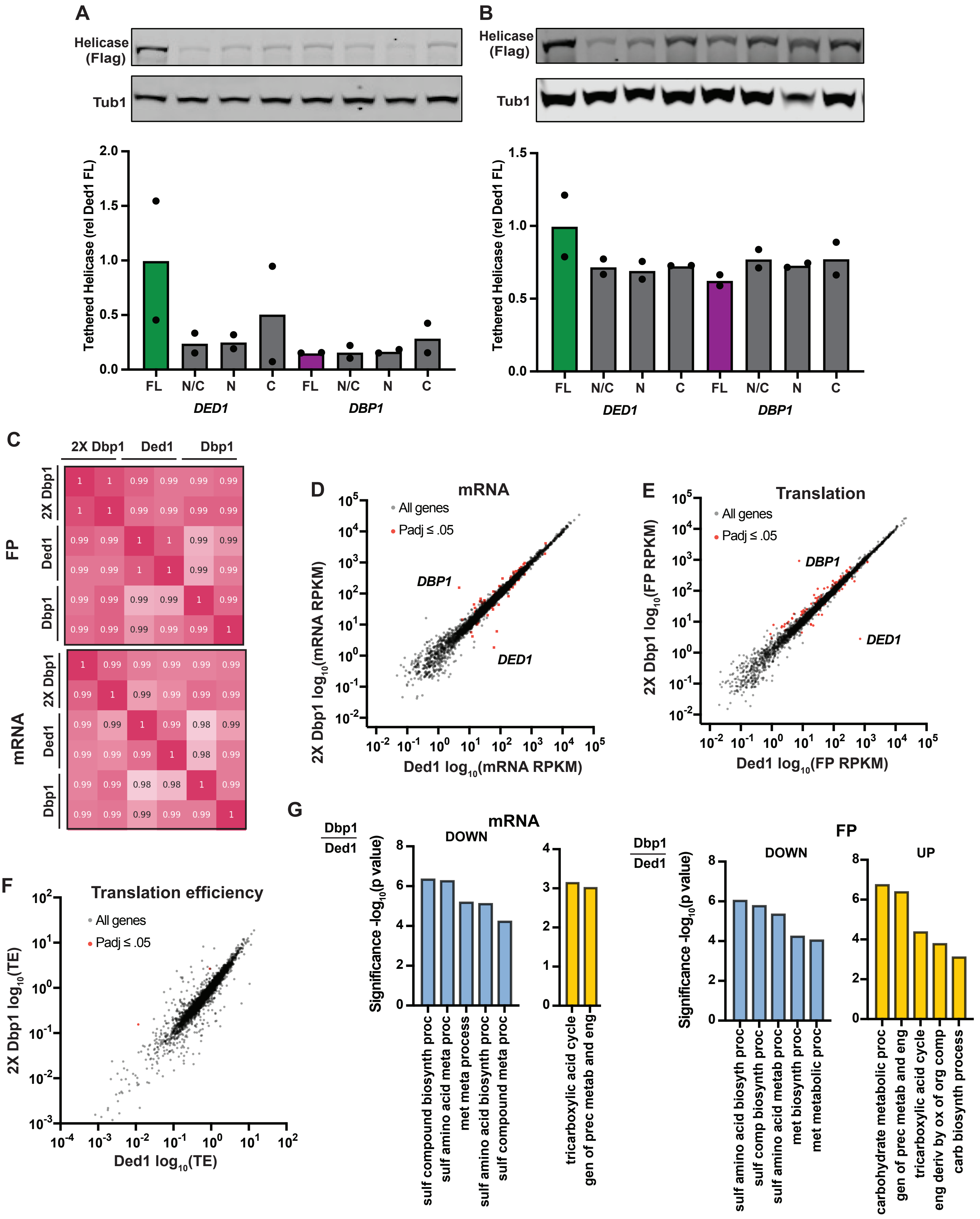
